# Electrostatic interactions define nacubactam potency against OXA-48-like β-lactamases

**DOI:** 10.1101/2024.10.30.621041

**Authors:** Joseph F. Hoff, Kirsty E. Goudar, Michael Beer, Philip Hinchliffe, John M. Shaw, Catherine L. Tooke, Yuiko Takebayashi, Adrian J. Mulholland, Christopher J. Schofield, James Spencer

## Abstract

Carbapenemases, β-lactamases hydrolysing carbapenem antibiotics, severely challenge treatment of multi-drug resistant bacterial infections. OXA-48 is among the most widely disseminated carbapenemases in *Enterobacterales*, consequently new treatment options for OXA-48 producers are urgently required. Development of diazabicyclooctane (DBO) inhibitors to overcome β-lactamase-mediated resistance is one attractive option, due to their efficacy against a wide range of β-lactamases. The DBO avibactam is already licensed for clinical use, with the related compound nacubactam currently in phase III trials for carbapenem-resistant infections. Here we investigate the activities of avibactam and nacubactam towards OXA-48 and two variants, OXA-163 and OXA-405, that contain deletions in the β5 - β6 loop adjacent to the active site and show modified activity towards different β-lactam classes. Compared to avibactam, nacubactam is c. 80-fold less potent towards OXA-48, but this difference is reduced in OXA-163 and OXA-405. Crystal structures of the respective avibactam and nacubactam complexes, and molecular dynamics simulations based upon these, reveal residue Arg214 on the OXA-48 β5 - β6 active-site loop to be electrostatically repelled by nacubactam, but not avibactam, binding. This increases flexibility of the OXA-48 β5 – β6 loop, as well as neighbouring active site loops, in simulations of the OXA-48:nacubactam, compared to the avibactam, complex. Such effects are not observed in simulations of the respective complexes of OXA-163 and OXA-405, which lack Arg214. These data indicate that interactions with Arg214 can determine DBO potency towards OXA-48-like enzymes, and suggest that sequence variation in this β-lactamase family affects reactivity towards inhibitors as well as β-lactam substrates.

## Introduction

Antimicrobial resistance (AMR) is a major and increasing threat to global public health, with 8.2 million associated annual deaths predicted by 2050^1^. The β-lactams (penicillins and related agents) are the most commonly prescribed antibiotic class^2,3^. In Gram negative bacteria, the dominant β-lactam resistance mechanism is expression of β-lactamase enzymes^4^ that hydrolyse the four-membered β-lactam ring (Fig. 1), to render β-lactams inactive towards their cellular targets, the penicillin-binding proteins (PBPs). β-Lactamases are divided, on the basis of sequence and structure, into four mechanistically distinct classes (A to D) of which classes A, C and D are serine β-lactamases (SBLs) and class B zinc-dependent metallo-β-lactamases (MBLs)^5^. OXA-48 is a class D SBL that uses a catalytic serine nucleophile (Ser70) to break down β-lactam substrates, with a proximal post-translationally carbamylated lysine (Lys73) proposed to act as the general base for both the acylation and deacylation steps of the hydrolysis reaction^6^.

**Figure 1:**
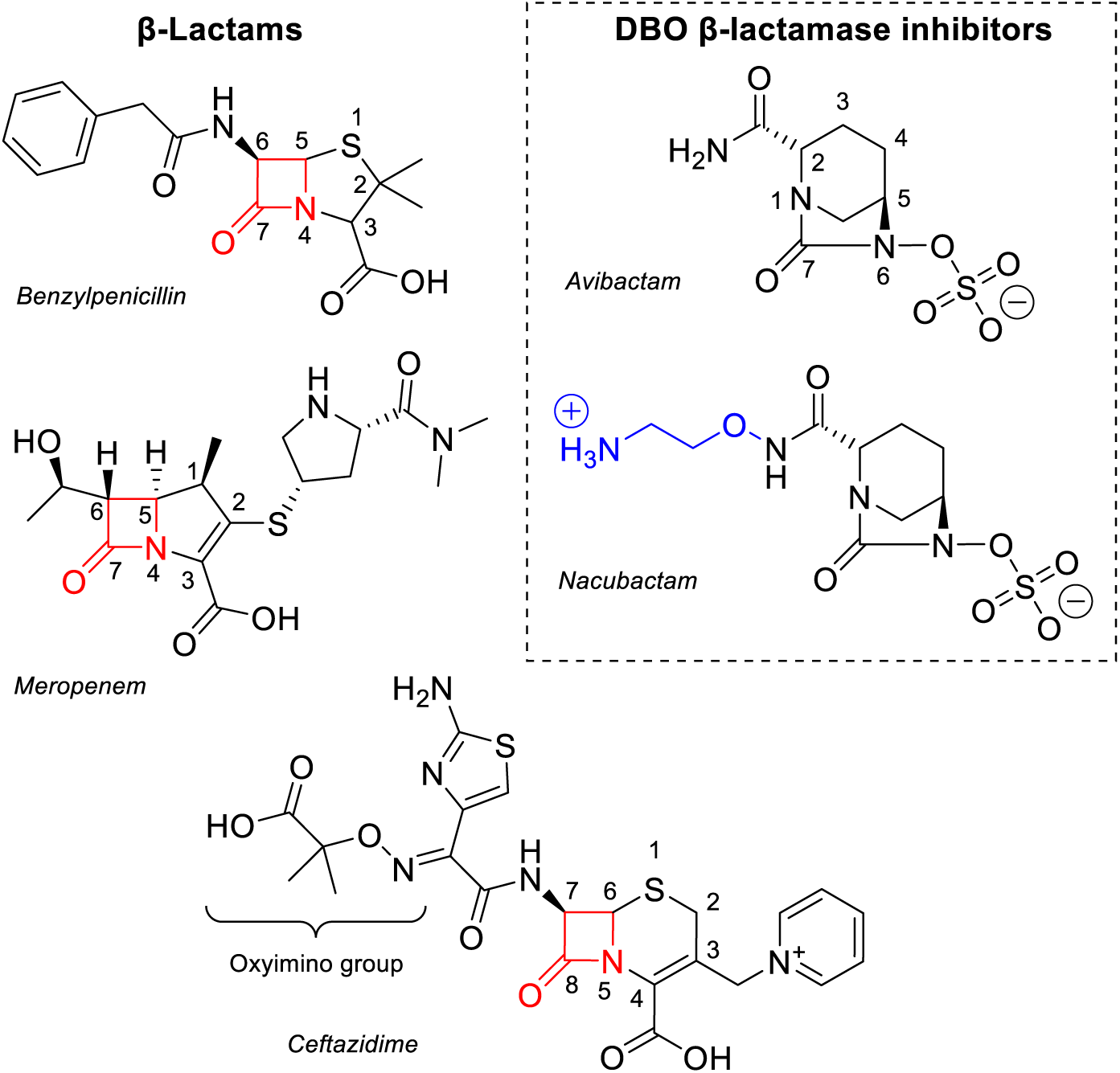
β-Lactam antibiotics and DBO β-lactamase inhibitors. Representative penicillin (benzylpenicillin), carbapenem (meropenem) and cephalosporin (ceftazidime) β-lactam antibiotics, with the β-lactam ring highlighted in red. DBO BLIs used in this work are shown on the right, with the additional 1-aminoethoxy group of nacubactam highlighted in blue. The core rings of avibactam and the β-lactams are numbered.

OXA-48 is of particular clinical importance due to its prevalence in the *Enterobacterales* order of bacteria, which includes two (*Escherichia coli* and *Klebsiella pneumoniae*) of the three pathogens most commonly associated with AMR-related deaths globally in 2019^7^. Worryingly, OXA-48 production causes failure of carbapenems, the class of β-lactam antibiotics reserved for the most serious bacterial infections. Due to its widespread dissemination, OXA-48 is regarded as one of the five most important acquired β-lactamases responsible for carbapenem resistance^8,9^. Carbapenem resistant *Enterobacterales* are classed by the World Health Organization as Critical Priority pathogens for global public health^10^.

To circumvent β-lactamase-mediated antibiotic resistance, β-lactamase inhibitors (BLIs) have been developed for co-administration with β-lactams^4^. The diazabicyclooctanes (DBOs) are one group of BLIs that have proven successful in the clinic, with avibactam currently used in combination with the expanded-spectrum oxyiminocephalosporin ceftazidime to treat multi-drug resistant bacterial infections (Fig. 1)^11^. DBOs inhibit SBLs through covalent attachment to the nucleophilic serine by a reversible carbamoylation reaction, and evade hydrolysis by recyclising to release the intact DBO (Fig. 2)^12,13^.

**Figure 2:**
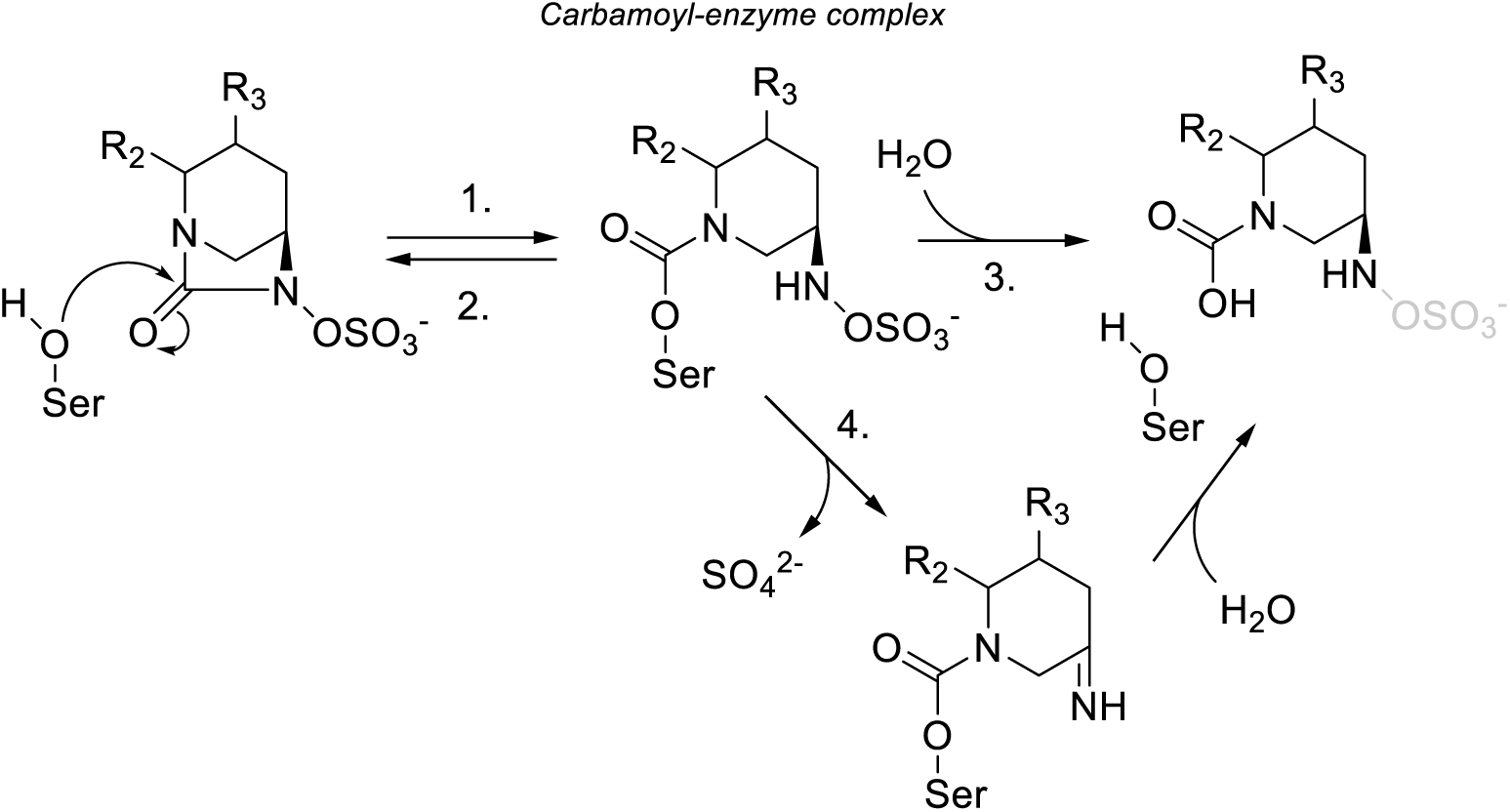
DBO inhibition and turnover by serine β-lactamases. DBO compounds inhibit SBLs by forming a covalent link with the catalytic serine via a (1.) carbamoylation reaction, analogous to acylation by β-lactam substrates. This results in formation of a carbamoyl-enzyme complex which has three proposed mechanistic fates: (2.) decarbamoylation to reform the intact DBO species, (3.) direct hydrolysis of the carbamoyl-enzyme complex and (4.) indirect hydrolysis via desulphation of the DBO-derived covalent intermediate^12–16^.

DBO inhibitors are of particular interest in the context of OXA-48, as this enzyme is generally poorly inhibited by more “classical” β-lactam-based inhibitors (tazobactam, clavulanic acid, sulbactam) but more effectively by avibactam^6,17^. In all currently available crystal structures of OXA-48 complexes, avibactam shows a very similar binding mode, and appears to induce decarbamylation of the catalytic Lys73, even under basic conditions, which has been attributed to the potency of avibactam inhibition towards class D carbapenemases^6,14,15,18^. However, Fröhlich *et al.*^11^ demonstrated that resistance to the ceftazidime:avibactam combination could be achieved in the laboratory through just two amino acid changes in the OXA-48 active site, highlighting the potential for changes in susceptibility to DBO combinations to emerge in clinical OXA-48 variants, and the need for continued inhibitor development to guard against such a possibility.

Following the development and introduction of avibactam^13^, numerous additional DBO inhibitors have been investigated, that largely differ in the composition of the C2 substituent group on the core DBO scaffold^19–21^. One such inhibitor, nacubactam, differs from avibactam by the addition of a 1-aminoethoxy group on the C2 group (Fig. 1), and is currently in phase III clinical trials in combination with β-lactams cefepime or aztreonam for the treatment of complicated urinary tract infections or uncomplicated pyelonephritis (ClinicalTrials.gov identifier NCT05887908)^22^. We have investigated activity of nacubactam, in comparison with avibactam, towards OXA-48 and two clinical variants, OXA-163 and OXA-405, that contain four amino acid deletions within their β5 – β6 active site loops and show reduced carbapenemase activity but enhanced hydrolysis of ceftazidime and other oxyiminocephalosporins. A combination of *in vitro* inhibition assays, X-ray crystal structures of enzyme:DBO complexes, and molecular dynamics simulations based upon these leads us to conclude that electrostatic repulsion between the positively charged N atom of the nacubactam C2 substituent and the side-chain of Arg214 on the β5 – β6 loop reduces potency towards OXA-48, compared to avibactam. These effects are less pronounced in the OXA-163 and OXA-405 variants, which lack Arg214. These data indicate that the effects of variation between OXA-48-like β-lactamases extend to interactions with DBO inhibitors, as well as classes of β-lactam substrates^23–25^.

## Results

### DBO interactions with the OXA-48 and its β5 – β6 loop variants, OXA-163 and OXA- 405

In this work, we sought to investigate the effects of DBO inhibitors upon naturally occurring OXA-48 variants differing in their activity towards expanded-spectrum oxyiminocephalosporin (e.g. ceftazidime) and carbapenem substrates. To understand the effect of β5 – β6 loop composition on DBO inhibition, we undertook *in vitro* inhibition experiments challenging OXA-48 and its naturally occurring variants, OXA- 163 and OXA-405, with the DBO inhibitors avibactam and nacubactam, which is distinguished from avibactam by an extended C2 substituent bearing a 1-aminoethoxy group. Both OXA-163 and OXA-405 have four amino acid deletions within the β5 – β6 loop, including of residue Arg214. OXA-163 also has a single amino acid substitution (Ser212Asp) adjacent to the deleted region. Compared to avibactam, nacubactam is a weaker inhibitor of OXA-48, with respective IC_50_ values of 19.91 µM and 0.26 µM, indicating a 77-fold difference in inhibition potency (Table 1). Although avibactam remains more potent than nacubactam towards both OXA-163 (IC_50_ values 0.20 µM and 1.23 µM, respectively) and OXA-405 (IC_50_ values 0.99 µM and 10.6 µM, respectively) for both variants there are notable increases in nacubactam inhibition potency compared to OXA-48 (16.2-fold for OXA-163, 1.9-fold for OXA-405). Moreover, the differences in inhibition potencies between avibactam and nacubactam are also reduced for the two variants (6.2- and 10.7-fold for OXA-163 and OXA-405, respectively). Therefore, the extended β5 – β6 loop of OXA-48 appears deleterious for the inhibition potency of nacubactam, compared to both OXA-163 and OXA-405.

**Table 1:**
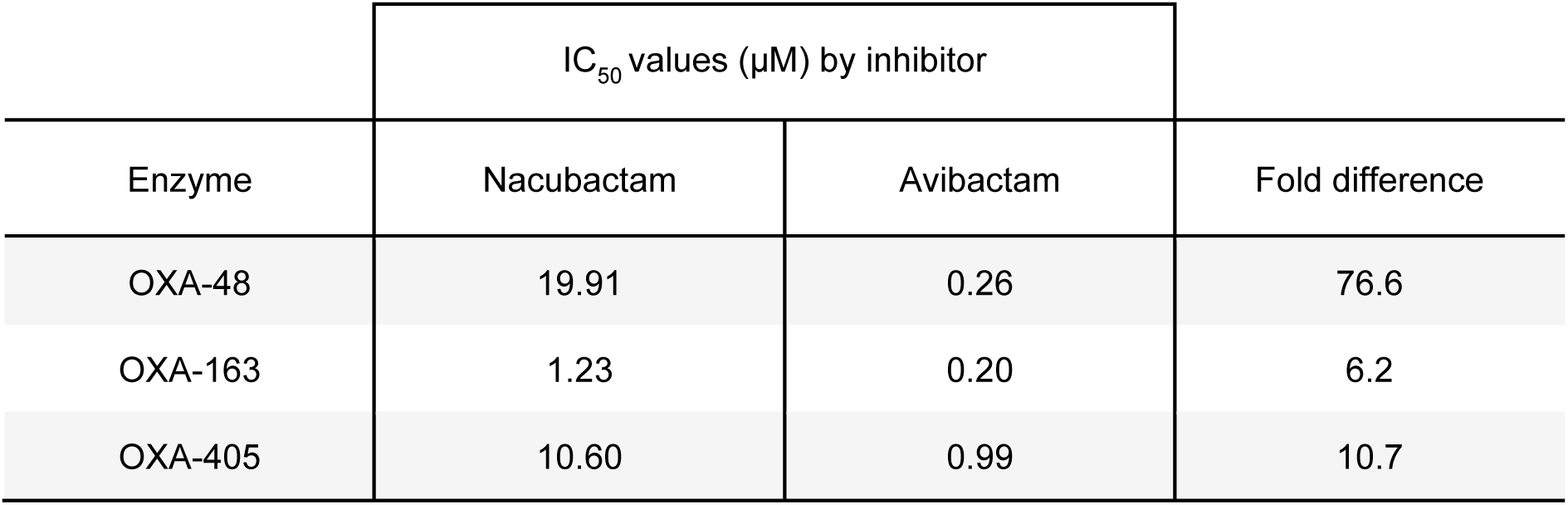
Inhibitory values (IC_50_) for nacubactam and avibactam against OXA-48- like β-lactamases.

However, minimum inhibitory concentration (MIC) values show nacubactam can better potentiate β-lactams ertapenem and ceftazidime against laboratory and control *E. coli* strains expressing these enzymes (Table S1). In fact, nacubactam reduces MICs at least 4-fold more greatly when combined with ertapenem and ceftazidime against *E. coli* expressing OXA-48, compared to avibactam. In contrast, nacubactam when combined with meropenem gives a 4-fold higher MIC value compared to meropenem with avibactam against OXA-48-producing *E. coli*, whereas more similar MICs are observed for the meropenem-avibactam combination against *E. coli* expressing OXA- 163 and OXA-405, corroborating with IC_50_ observations.

### Crystal structure of OXA-48:nacubactam carbamoyl-enzyme complex

To understand the basis for these differences in DBO inhibition potency towards OXA- 48 and the two variants in more detail, we sought to determine crystal structures for the respective complexes, comparing as appropriate with pre-existing structures. The crystal structure of nacubactam bound to OXA-48 was solved at 1.51 Å resolution with the enzyme assembled as a homodimer and a chloride ion positioned at the dimer interface, as characterised previously (Fig. 3A, Table S2)^26^. Continuous positive *F*_o_-*F*_c_ electron density between the ligand and Ser70 indicates the presence of a covalently bound nacubactam-derived carbamoyl-enzyme complex (Fig. 3B). When overlaid upon the structures of uncomplexed OXA-48 (1.38 Å resolution, Table S2) and OXA- 48 bound to avibactam (PDB 4S2K^14^, 2.10 Å), all three structures have very similar backbones, as highlighted by the low Cα RMSD values when aligned (0.41 Å and 0.53 Å for superposition of the OXA-48:nacubactam complex on the uncomplexed and avibactam-bound structures, respectively, as calculated in PyMOL). The avibactam and nacubactam carbamoyl-enzyme complexes show a similar binding modes to OXA-48, with the C7 carbonyl group positioned within the oxyanion hole formed by the backbone amides of residues Ser70 and Tyr211, as seen in β-lactam-derived acyl-enzyme complexes of OXA-48-like enzymes (Fig. 3C)^6^. Both complexes also show the DBO sulphonate group pointing towards the side chains of residues Arg250 and Thr209, with the side chain hydroxyl of Ser118 within hydrogen bonding distance of the DBO N6. Thus, binding of the DBO core is facilitated by complementary electrostatic and hydrogen-bonding interactions, whilst the respective C2 substituents point out of the active site towards the OXA-48 β5 - β6 (residues 212 - 219) and Ω- (residues 144 - 163) loops.

**Figure 3:**
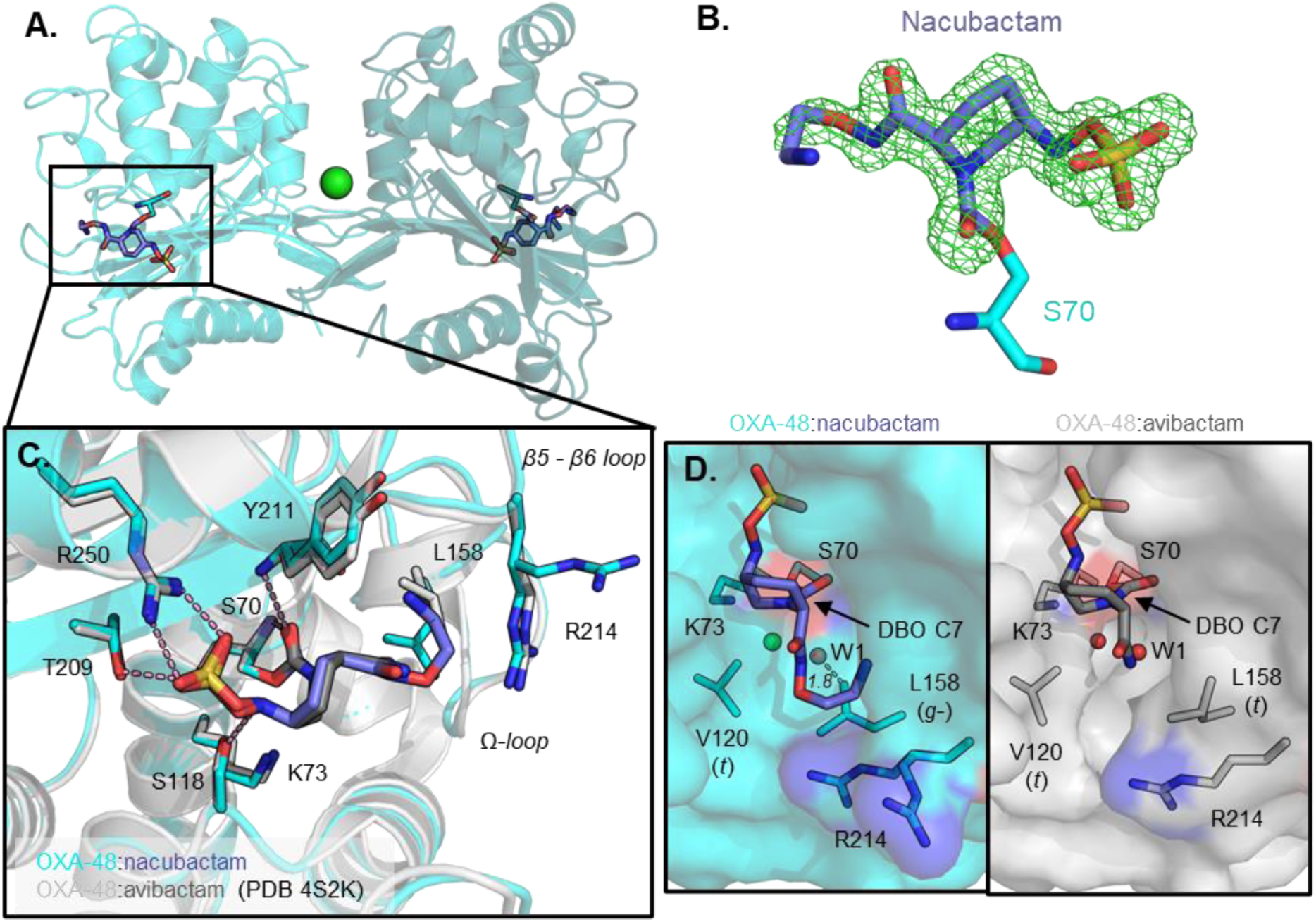
Crystal structure of OXA-48 bound to nacubactam. (A) Crystal structure of OXA-48 carbamoyl-enzyme complex with nacubactam. Different shades of cyan represent different chains in the homodimer, chloride ion at the dimer interface is shown as a green sphere. (B) Nacubactam-derived carbamoyl-enzyme complex, mesh shows F_o_-F_c_ electron density from refinement carried out in the absence of ligand, contoured at 3σ. (C) Close-up of the chain B active site (cyan), overlaid with a previously determined structure of OXA-48 bound to avibactam (PDB 4S2K^14^, grey). (D) Surface views of the deacylating water channel in DBO-bound OXA-48 complexes, water molecules and chloride ions are represented as red and green spheres, respectively. Leu158 χ_1_ (N-Cα-Cβ-Cγ) and Val120 χ_1_ (N-Cα-Cβ-Cγ1) rotamers are highlighted in brackets; dashed line shows distance between the dual-occupancy water molecule (W1) and Leu158 Cδ2.

One noticeable difference between the active site arrangements of these complexes is the carbamylation status of Lys73, which is decarbamylated in both DBO-bound structures (determined at pH values of 8.8 and 7.5 for the nacubactam and avibactam complexes, respectively, Table S3), but remains carbamylated in the uncomplexed enzyme (structure determined at pH 7.5, Fig. S1, Table S3)^14^. An additional difference is in the positioning of the side chain of residue Leu158 in the active-site Ω-loop, which adopts a *g-* χ_1_ side-chain rotamer in the avibactam complex, whereas the *t* rotamer is observed in uncomplexed and nacubactam-bound OXA-48 (Fig. 3). It is likely that Leu158 adopts this conformation to avoid steric clashes with the bulkier C2 substituent of nacubactam, compared to avibactam. As Leu158 is part of the so-called ‘deacylating water-channel’, differences are observed in solvent access to the hydrophobic pocket in which Lys73 resides (Fig. 3D)^6,27,28^. Surface views suggest that when Leu158 is in the *t* rotamer, as in the avibactam complex, this channel is open, as evidenced by presence of a water (W1) proximal to the C7 carbonyl of the carbamoyl-enzyme complex. In contrast, when Leu158 adopts the *g-* rotamer, observed when nacubactam is bound, the channel is closed, and the adjacent water is modelled in dual occupancy with the Leu158 side-chain due to steric clashes (Leu158 Cδ2 is ∼1.8 Å away from the position of this water molecule) (Fig. 3D).

In the active site of chain B, Arg214 of the Ω-loop adopts a dual conformation when OXA-48 is bound to nacubactam. In both conformations, the Arg214 side chain is poorly resolved in the final 2*F*_o_-*F*_c_ electron density map (Fig. S2). Thus, it is possible that the side-chain of Arg214 and the C2 tail nitrogen of nacubactam are electrostatically repelled *in crystallo* as a result of their clashing positive charges. Chain A of the nacubactam complex, in contrast, does not show this effect, most likely due to a crystal packing artefact whereby residue Glu132 from an adjacent molecule not in the asymmetric unit can electrostatically interact with Arg214, thus holding it in place. Consequently, the C2 tail group of nacubactam adopts a dual conformation in the chain A active site, presumably as a result of electrostatic repulsion by the Arg214 side-chain. Of note, in these experiments OXA-48 was purified and crystallised under basic conditions (purification buffer pH 8.4, crystallisation solution pH 8.8) but it is very likely that both Arg214 and the nitrogen atom of the nacubactam C2 substituent are protonated at physiological pH. Indeed, the predicted pKa of Arg214 is 12.2 (PropKa webserver^29^) and that of the nacubactam C2 aminoethoxy nitrogen 8.7 - 9.1 (AIMNet2/MolGpka webservers^30,31^, Fig. S3).

### MM MD simulations of DBO-derived carbamoyl-enzyme complexes

To investigate the possibility that these differences between avibactam and nacubactam binding affect the dynamic behaviour of OXA-48, MM MD was next used to simulate uncomplexed, avibactam- and nacubactam-bound OXA-48. Root mean-squared deviation (RMSD) plots indicate that all simulations were stable over total trajectories of 1.5 µs (Fig. S4). Pairwise differences in average root mean-square fluctuation (ΔRMSF) for each residue, across both active sites of the OXA-48 dimer, were then calculated between the different simulations (Fig. 4). When comparing simulations of the two DBO-bound complexes, three regions of the OXA-48 sequence, the Ω- (residues 144 - 163), β5 – β6 (212 - 219) and β7 – α10 (240 - 247) loops show noticeable changes in ΔRMSF (Fig. 4A). When mapped onto the structure of OXA-48, these regions of difference are centred around Arg214, which has an average ΔRMSF of +0.41 Å. This indicates an increase in mobility of this residue in the nacubactam, compared to the avibactam complex, consistent with our crystallographic evidence (above) of electrostatic clashes between the nacubactam C2 substituent and the β5 – β6 loop (Fig. 3C). Indeed, measuring the distance between Arg214 (Cζ) and the C2 amide nitrogen atoms of each DBO inhibitor over the respective simulation trajectories reveals that this distance is generally longer, with Arg214 adopting a more variable position, in the nacubactam-, compared to the avibactam-bound, complex (Fig. S5).

**Figure 4:**
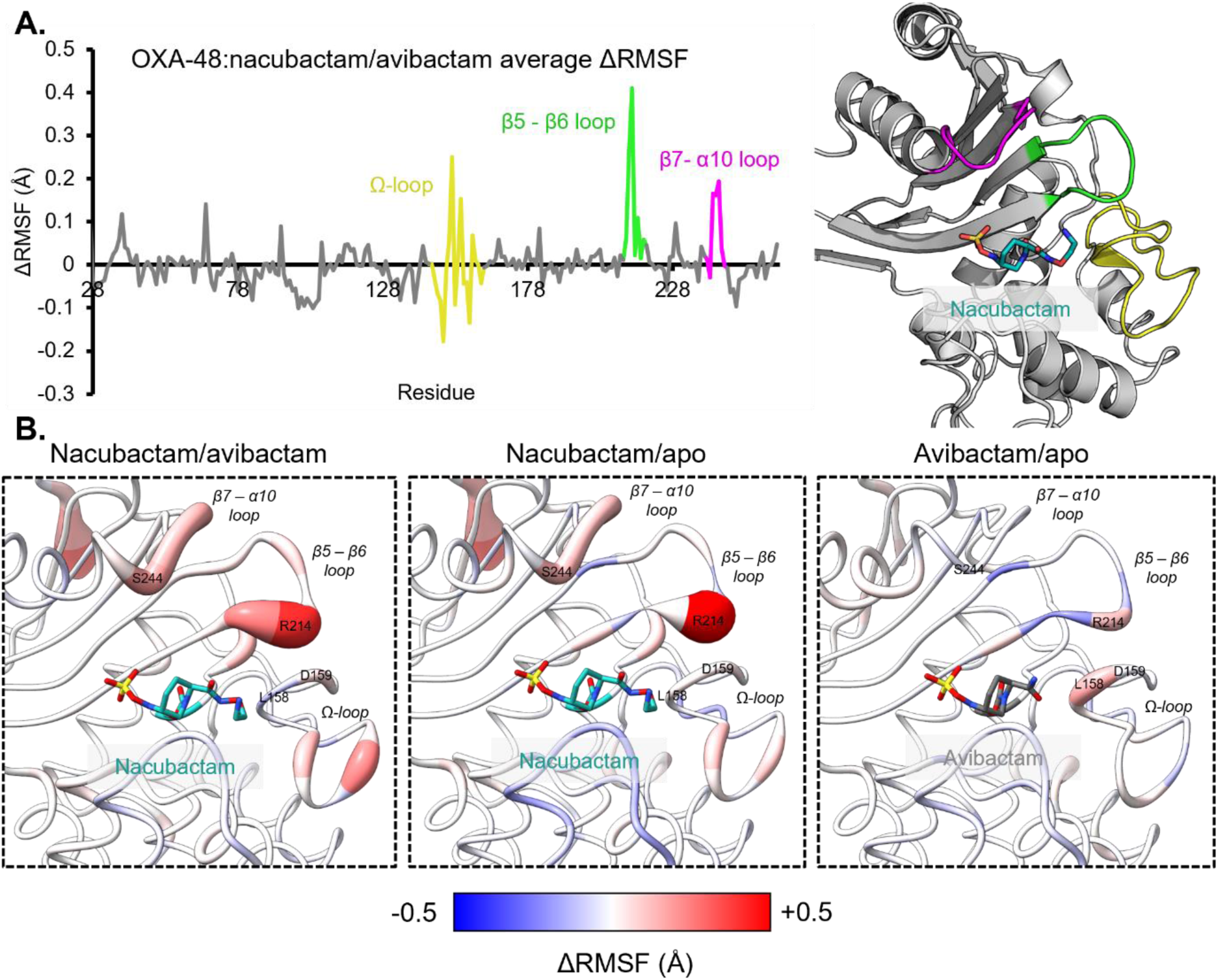
RMSF analysis of MM MD simulations of OXA-48:DBO carbamoyl-enzyme complexes. (A) Difference in residue RMSF (ΔRMSF) between simulations of DBO-bound OXA-48, averaged across both chains of the protein homodimer, plotted against enzyme primary sequence. More positive values indicate greater fluctuation in nacubactam-, compared to avibactam-bound, OXA-48. The OXA-48 Ω- (yellow, residues 144 - 163), β5 – β6 (green, residues 212 - 219) and β7 – α10 (pink, residues 240 - 247) loops are highlighted on the graph and OXA-48:nacubactam crystal structure. (B) Comparisons of ΔRMSF values, coloured according to scale below, between DBO-bound and uncomplexed OXA-48 simulations, mapped onto the crystal structures of OXA-48 bound to nacubactam and avibactam. Thicker and more red backbones indicate more positive ΔRMSF values.

In addition, further simulations of the OXA-48:nacubactam complex, in which the C2 aminoethoxy nitrogen was deprotonated (Figures S4, S5), do not show the same degree of repulsion of Arg214, suggesting that the observed displacement is largely due to electrostatic effects.

Arg214 displacement by nacubactam results in an increase in flexibility across adjacent loops in the OXA-48 active site, most notably within the β5 - β6 loop itself and, by disruption of a salt bridge with Asp159, across the Ω-loop (Fig. 4B). The β7 – α10 loop, which does not directly interact with bound DBO inhibitors in carbamoyl-enzyme complexes, also appears to be destabilised by increased movement of the β5 – β6 loop. When compared to simulations of uncomplexed OXA-48, the nacubactam-bound complex shows similar RMSF differences to those obtained from comparison with the avibactam complex. In contrast, such differences are less prominent when the avibactam-bound structure is compared with that of uncomplexed OXA-48, although the Arg214 RMSF increases slightly in the former. However, this does not noticeably affect the stability of neighbouring loops in the same way as was observed for nacubactam-bound, compared to uncomplexed, OXA-48.

In contrast, Leu158, on the Ω-loop, appears more flexible when avibactam is bound, compared to uncomplexed and nacubactam-bound OXA-48. In simulations of nacubactam-bound OXA-48 Leu158 predominantly adopts the *g-* χ_1_ rotamer, as observed in the crystal structure (Fig. S6A). In contrast, in the avibactam-bound or uncomplexed structures the *t* rotamer is preferred. Conformation of the Leu158 side-chain is therefore specifically limited in the nacubactam-bound complex, as a result of steric restrictions imposed by its larger C2 substituent, compared to that of avibactam, as observed *in crystallo* (Fig. 3D). In contrast, Val120, the other residue contributing to the OXA-48 deacylating water channel^27,28^, consistently adopts the *t* rotamer, as observed in the crystal structures, in all simulations of the uncomplexed and DBO- bound enzyme (Fig. S6B).

### DBO complexes with OXA-48 variants OXA-163 and OXA-405

Crystal structures were next obtained for the OXA-48 variants OXA-163 and OXA-405, as uncomplexed enzymes and in the nacubactam- and avibactam-bound forms (Table S2, Fig. S7). Both enzymes assemble as homodimers in the asymmetric unit and, consistent with their high overall sequence identities, adopt very similar global structures and active site arrangements to OXA-48. The exception is the Ser70 nucleophile, which is in dual conformation in one of the two chains in the crystallographic dimers of uncomplexed OXA-163 and OXA-405 (Fig. S8A, B). The respective protein backbones only differ at the β5 – β6 loop, which is truncated in the cases of OXA-163 and OXA-405, compared to OXA-48 (Fig. S8C). Part of the OXA- 405 β7 – α10 loop, which neighbours the β5 – β6 loop, could not be modelled due to weak electron density, indicating that this loop is more mobile in uncomplexed OXA- 405, compared to the other enzymes. Lys73 is fully decarbamylated in all DBO-bound structures except the OXA-405:avibactam (both chains) and OXA-163:avibactam (only in chain A) complexes, where Lys73 is modelled as partially carbamylated (Table S4A), in dual-occupancy with a free lysine side-chain that is associated with either a water molecule or chloride ion (Fig. 5A).

**Figure 5:**
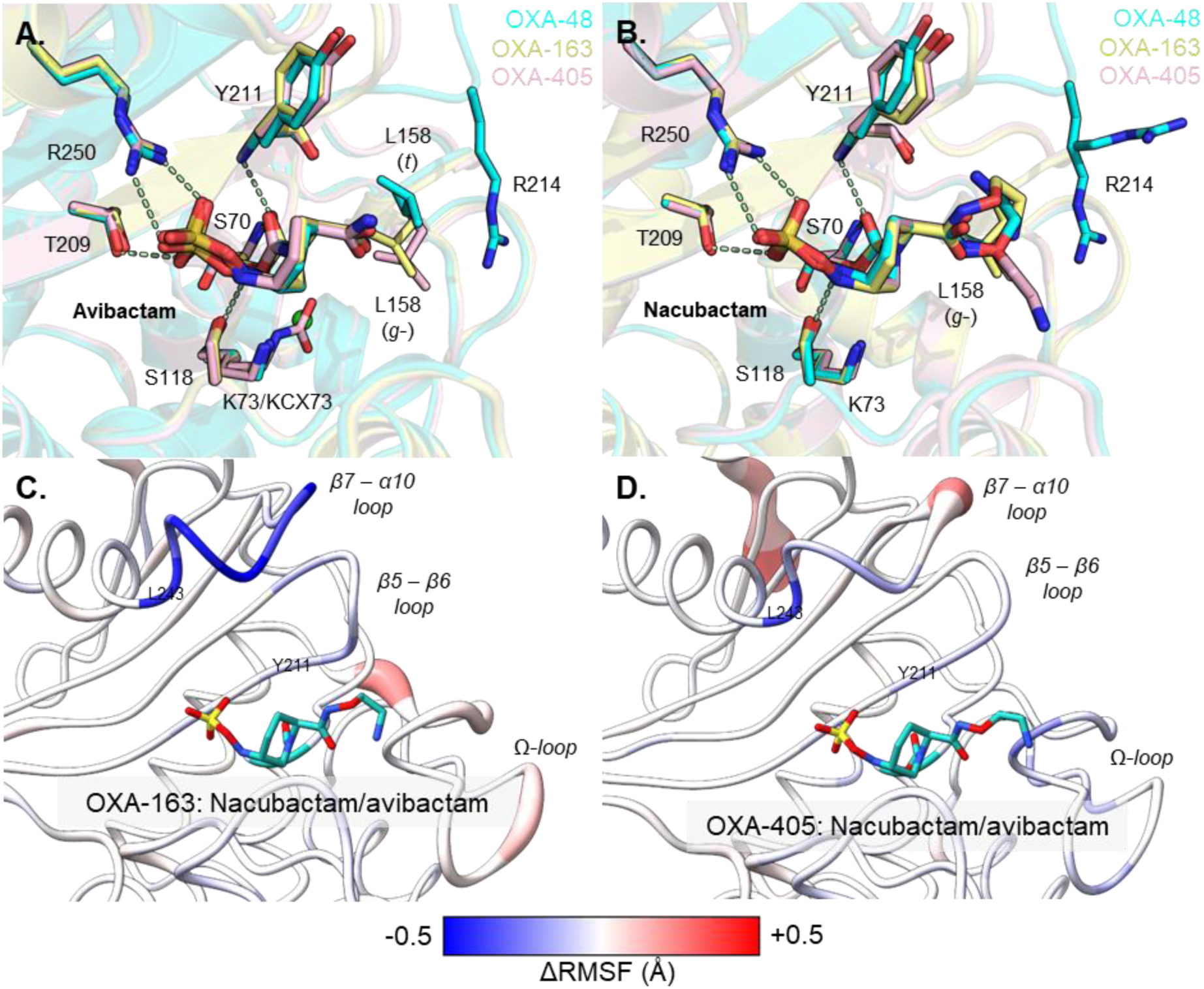
Nacubactam and avibactam inhibition of naturally occurring OXA-48 variants OXA-163 and OXA-405. (A) and (B) Active site overlays of OXA-48 (turquoise), OXA-163 (yellow) and OXA-405 (pink) bound to (A) avibactam and (B) nacubactam. Residues are shown as sticks, hydrogen bonding interactions in OXA- 48:DBO structures as dashed lines and chloride ions as green spheres. Residues are numbered according to OXA-48, Leu158 χ_1_ rotamer form is shown in brackets. (C) and (D) ΔRMSF analysis of MM MD simulations of (C) OXA-163 and (D) OXA-405 complexes with nacubactam and avibactam, mapped onto crystal structures of the respective nacubactam complexes.

In chain B of the OXA-405:nacubactam complex Tyr211 adopts a dual conformation (Fig. 5B). The presence of an alternative side-chain rotamer coincides with poor electron density for the β7 – α10 loop, as also seen in uncomplexed OXA-405 (Fig. S9). Furthermore, Leu243 of the β7 – α10 loop also adopts multiple conformations to avoid steric clashes with the Tyr211 side chain. In contrast, in the active site of chain A in the OXA-405:nacubactam complex, residues Tyr211 and Leu243 are each in a single conformation and the β7 – α10 loop can be modelled in its entirety.

In the respective carbamoyl-enzyme complexes with OXA-405 and OXA-163 nacubactam and avibactam adopt similar binding modes to those observed for the OXA-48 parent enzyme (Fig. 5A, B), characterised by complementary electrostatic interactions between the DBO sulphonate group and Arg250 (OXA-48 numbering) and positioning of their C2 tail groups close to the β5 – β6 loop. The C2 substituent of avibactam is identically positioned in all three enzymes, whereas the additional 1- aminoethoxy group of nacubactam appears more variably located across these structures. Indeed, the unbiased electron density for this extended substituent of nacubactam is less well defined compared to that for the C2 amide component common to the two DBOs, suggesting that in all three enzymes the nacubactam C2 substituent is much more flexible than is the case for avibactam (Fig. S7).

Residue Leu158 appears to adopt an alternative rotamer (*g-*) when avibactam is bound to OXA-163 (chain B) and OXA-405, compared to OXA-48 or chain A of the OXA-163:avibactam complex where the *t* rotamer is observed. In contrast, when nacubactam is bound, Leu158 is in the *g-* rotamer for OXA-48 and OXA-163, whilst in the OXA-405:nacubactam complex both rotamers are observed across the two active sites. Thus, the relationship between the conformation of Leu158, the composition of the β5 – β6 loop, and susceptibility to DBO inhibition appears to be structurally complex in the context of OXA-48-like enzymes.

### MM MD simulations of DBO-bound OXA-163 and OXA-405 complexes

MM MD simulations of uncomplexed, avibactam- and nacubactam-bound OXA-163 and OXA-405 were then run, using the same protocol as for OXA-48. RMSD plots (Fig. S4) indicate all simulations to be stable over their respective trajectories. ΔRMSF analysis reveals no noticeable differences in the flexibility of the β5 - β6 loop between simulations of avibactam- and nacubactam-bound OXA-163 and OXA-405 (Fig. 5C, D). The Ω-loop appears slightly more flexible in nacubactam-, compared to avibactam-bound, OXA-163, although the difference is less marked than is the case for OXA-48 (Fig. 4B). The β7 - α10 loop, in contrast, is much more stable in both of the nacubactam, compared to the avibactam, complexes. This corresponds with a reduction in mobility of Tyr211, suggesting that in OXA-163 and OXA-405 nacubactam can stabilise this residue much more readily than does avibactam. This in turn may affect the dynamics of the β7 - α10 loop, consistent with our crystal structures (Fig. S9). Indeed, residue Leu243, which is shown to be dislodged by conformational changes of Tyr211 in the crystal structure of OXA-405 bound to nacubactam, is much more stable in both cases.

Ser118 is suggested to play an important role in DBO turnover by SBLs^14,15^. Proposed reaction mechanisms for class D SBLs suggest that Ser118 can deprotonate the N6 atom of the DBO carbamoyl-enzyme to promote intramolecular recyclisation, and may also contribute to activation of the decarbamoylating water for hydrolysis^6,12^. Therefore, the distances between the DBO N6 and Ser118 Oγ atoms were measured during our MM MD simulations, and the percentage of simulation frames where this distance is within an arbitrarily selected 3.5 Å cutoff calculated (Fig. 6). No major differences in these percentage values were observed between the two independently modelled active sites in the various dimeric complexes, with the largest difference 11.2% between chains A and B in the simulation of avibactam-bound OXA-163. However, the simulations revealed that, in complexes of OXA-48, OXA-163 and OXA- 405, the N6 of nacubactam was more frequently positioned close to the Ser118 side chain than was the case for avibactam. These differences, when averaged across both active site complexes, are greatest for OXA-163 (44.0%), followed by OXA-405 (34.4%), with OXA-48 showing the smallest difference in the Ser118:DBO N6 distance between the avibactam- and nacubactam-bound complexes (17.5%). However, in simulations of the OXA-48:nacubactam complex in which the N of the nacubactam C2 substituent is deprotonated, the distance between Ser118 and the DBO nitrogen N6 is consistently greater than in any of the other enzyme:DBO complexes studied here, with these atoms positioned within 3.5 Å of each other in only 10.1% of simulation frames. This analysis suggests that protonation of the nacubactam C2 tail group is required for positioning close to Ser118, and can consequently affect the propensity for recyclisation.

**Figure 6:**
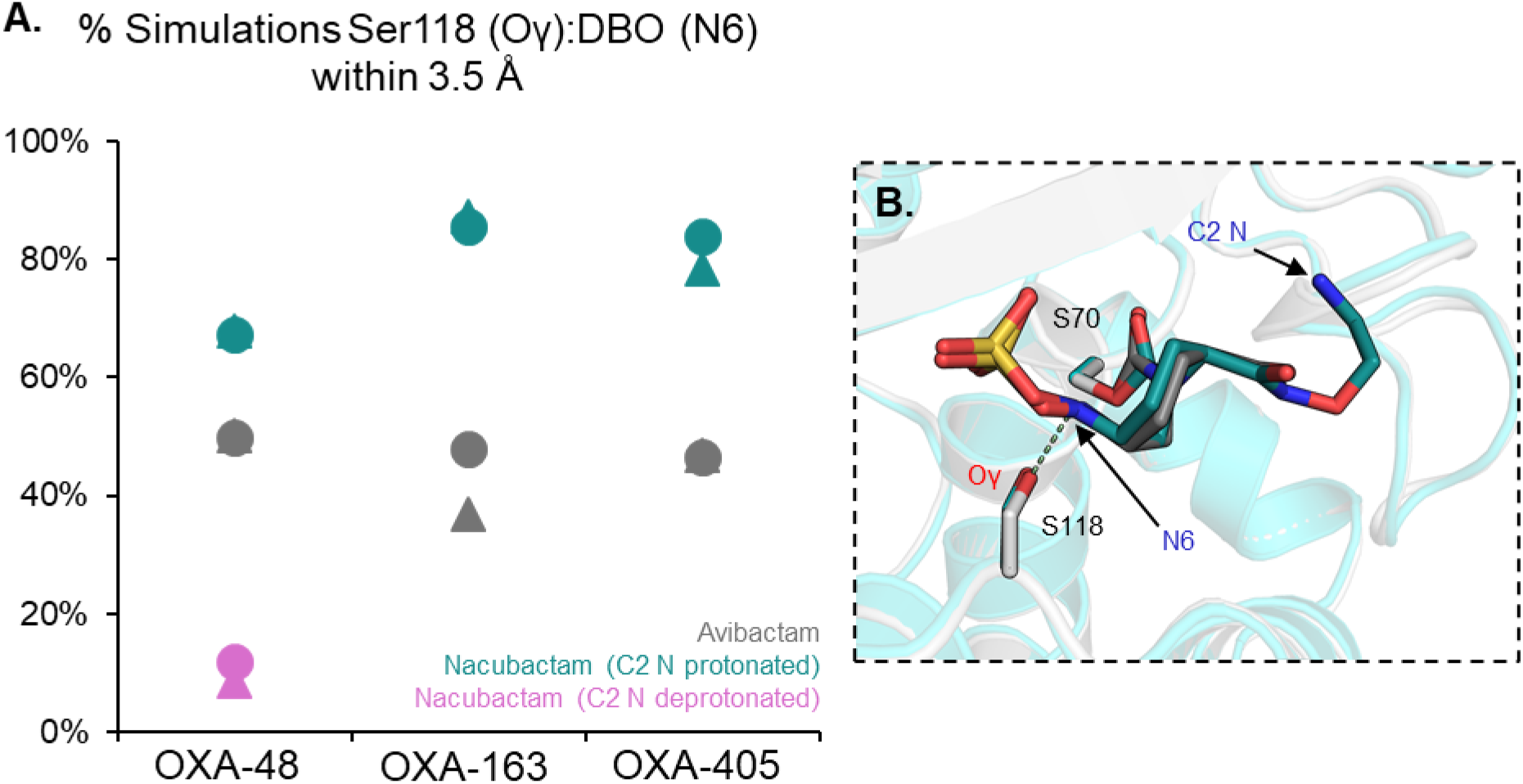
Analysis of distances between Ser118 Oγ and DBO N6 atoms in simulations of DBO carbamoyl complexes of OXA-48-related enzymes. (A) Proportion (%) of simulations where Ser118 (Oγ) and N6 of the DBO-derived carbamoyl-enzyme complex are within 3.5 Å of each other, for each OXA-48-like enzyme. Triangles represent chain A and circles chain B of individual DBO complexes. (B) Overlay of OXA-48 avibactam- (PDB 4S2K^14^, grey) and nacubactam-bound (turquoise) structures, with the Ser118:DBO distance represented by a pale green dashed line.

## Discussion

OXA-48 has become one of the most commonly detected carbapenemases in *Enterobacterales* globally, thus treatment options that circumvent β-lactam antibiotic resistance as a result of OXA-48-production are of clinical importance^8^. DBO-based BLIs are an established option, and various efforts have been made to optimise their activity, with multiple compounds having modified C2 substituents compared to avibactam, the original DBO now available in the clinic^4^. The degree of modification to the C2 substituent appears to correlate with intrinsic antimicrobial, as well as β- lactamase inhibitory, activity resulting from the ability to bind to both PBPs and SBLs. This gives rise to the possibility of a ‘dual-action’ agent that would not require combination with a β-lactam for treatment of antibiotic-resistant infections^32,33^. Indeed, nacubactam, which compared to avibactam has an extended 1-aminoethoxy group on its C2 substituent, has been shown to have some antimicrobial activity, attributed to its ability to inhibit PBP2 in *Enterobacterales*^22^.

Here, we have biochemically and structurally characterised inhibition by nacubactam, in comparison with avibactam, of OXA-48 and its naturally occurring variants OXA- 163 and OXA-405. IC_50_ values show nacubactam to be a substantially weaker inhibitor of OXA-48 than avibactam, whereas differences in the potencies of the two DBOs decrease in OXA-163 and OXA-405, which both have four amino acid deletions in the β5 – β6 active site loop. However, we observe potency differences in IC_50_ and MIC experiments. This could be due to the improved cell uptake of nacubactam and/or its propensity to inhibit PBP2 that may enhance β-lactam action, as seen by a reduction in MICs against non-β-lactamase-producing lab/control strains (Table S1)^22^.

We determined high-resolution crystal structures of uncomplexed, avibactam- and nacubactam-bound OXA-48, OXA-163 and OXA-405 to investigate how differences in β5 – β6 loop composition relate to DBO potency. This revealed conformational variability of this loop only in the OXA-48:nacubactam complex, as a result of electrostatic repulsion between the C2 tail nitrogen of the nacubactam 1-aminoethoxy group and the side chain of Arg214. Subsequent MM MD simulations of these structures reveal increased flexibility of the β5 – β6 loop in nacubactam-bound OXA- 48, likely due to DBO-mediated electrostatic displacement of Arg214, which consequently appears to propagate flexibility in neighbouring active site loops. Such an effect has also been shown in a structure of OXA-48 in complex with the oxyimino-cephalosporin antibiotic ceftazidime (PDB 6Q5F), where the extra bulk of the ceftazidime oxyimino group (Fig. 1) appears to disrupt the interaction between Arg214 and Asp159, which the authors suggest increases flexibility of the Ω-loop, as evidenced by a lack of electron density for this region in the crystal structure^11^. Therefore, it appears that Arg214 is an important determinant of OXA-48 active site conformational flexibility, and its displacement may begin to explain the weaker IC_50_ value of nacubactam towards OXA-48, compared to avibactam.

We note, however, that nacubactam potency is not fully restored to levels observed for avibactam when Arg214 is deleted, as in OXA-163 and OXA-405, suggesting that additional factors can affect DBO potency towards these enzymes. In fact, in MM MD simulations of all three enzymes, we see a greater propensity for nacubactam, compared to avibactam, to adopt a binding pose that may more readily facilitate DBO recyclisation, as indicated by the closer proximity of the DBO N6 nitrogen and the Ser118 side-chain oxygen. Thus, it is possible that, compared to avibactam, nacubactam is more readily recyclised by OXA-48-related enzymes, therefore reducing its inhibitory potency by reducing the lifetime of the carbamoyl-enzyme complex.

We also see differences between the various complexes of OXA-48 family members, in both crystal structures and in simulations, in the positioning of residue Leu158 in the deacylating water channel. It appears that steric considerations with respect to the extended C2 1-aminoethoxy substituent of nacubactam restrict the flexibility of Leu158, thus resulting in a closed deacylating water channel when OXA-48 is bound to nacubactam. This may affect solvent accessibility of the hydrophobic pocket in which Lys73 resides, in turn influencing the carbamylation status of Lys73 due to the requirement for water access to mediate lysine decarbamylation^12,34–36^. This has been observed in time-resolved crystallographic studies of avibactam carbamoylation in the class D SBL CDD-1, that show corresponding movement of Leu158 and release of carbon dioxide following Lys73 decarbamylation^27^. The consequence, for DBO susceptibility, of Leu158 side-chain position and Lys73 carbamylation status is likely to be complex due to the multiple DBO turnover pathways that can occur in class D SBLs (Fig. 2)^12^. Indeed, the variability of Leu158 rotamers in our crystal structures of DBO complexes with OXA-163 and OXA-405 highlights this.

In conclusion, this work suggests a structural relationship between DBO inhibitory potency and β5 - β6 active site loop composition in class D β-lactamases of the OXA- 48 family. Our findings thus suggest that, while variants differ in susceptibility to individual DBOs, rational design strategies involving optimising C2 substituents to make more favourable interactions with the active site is one route to more potent DBO inhibitors of OXA-48 family members.

## Methods

### Recombinant enzyme expression and purification

For crystallography and kinetic experiments, genes encoding the mature OXA-163^23– 261^ and OXA-405^23-265^ open reading frames were synthesised (Eurofins Genomics), amplified by PCR and cloned into the pOPINF T7 expression vector by recombination (InFusion, Takara) using primers listed in Table S5^37^. *E. coli* BL21-DE3 cells (Novagen) were transformed with pOPINF-OXA-48^(23-265)^ (previously cloned as described^38^), pOPINF-OXA-163^(23-261)^ and pOPINF-OXA-405^(23-261)^ by heat shock and grown in 2X YT broth supplemented with 50 µg.mL^-1^ carbenicillin (Fisher Scientific) for large scale growth (37 °C, 180 rpm shaking). When the culture reached an OD_600_ of 0.6 – 0.8, enzyme expression was induced with the addition of 0.5 mM IPTG (Calibre Scientific) and left overnight at 18 °C, 180 rpm shaking. Cells were then pelleted by centrifugation (6500 × *g*, 10 minutes) and resuspended in 50 mL of the respective purification buffer (50 mM Tris pH 8.4, 0.25 M NaCl, 20% Glycerol (OXA-48); 150 mM HEPES pH 8.0, 0.4 M NaCl (OXA-163); or 150 mM HEPES pH 6.5, 0.4 M NaCl (OXA-405)) supplemented with 10 mM imidazole, 1 tablet of cOmplete EDTA-free protease inhibitor cocktail (Roche), 1 µL Benzonase Endonuclease (Novagen) and 2 - 5 mg lysozyme (Sigma). All subsequent purification steps were completed at 4 °C or on ice. Cells were lysed by a cell disruptor (Constant Systems) at 25 kpsi and centrifuged at 100,000 × *g* for 1 hour. The soluble fraction was then added to 4 mL Ni-NTA beads (Qiagen) pre-washed in water, and incubated for 1 hour with rotation. The beads were washed once with 10 mL of the respective purification buffer supplemented with 10 mM imidazole, and once more with 10 mL purification buffer supplemented with 20 mM imidazole. Protein was then eluted and collected with the addition of 10 mL of the respective purification buffer supplemented with 300 mM imidazole. NaCl was omitted from the elution step for OXA-163 and OXA-405. The eluent was buffer-exchanged to reduce the imidazole concentration to less than 50 mM using a 10 kDa cutoff Vivaspin Centrifugal Concentrator (Sartorius), and subsequently incubated overnight with 2 mg recombinant 6His-tagged 3C protease to cleave the enzyme purification tag. The mixture was run through a second 10 mL Ni-NTA (nitrilotriacetic acid) column to remove 3C protease and cleaved hexhistidine tags, and the eluent from this column collected, concentrated to 5 mL and loaded onto a 120 mL HiLoad 16/600 Superdex 75 size-exclusion column (Cytiva) equilibrated with OXA-48 purification buffer, or 0.1 M sodium phosphate buffer at pH 8.0 or pH 6.5 for OXA-163 or OXA-405, respectively. Peak fractions were collected (purity determined by SDS-PAGE^39^), pooled and concentrated to 11.5 mg.mL^-1^ (OXA-48), 7.7 mg.mL^-1^ (OXA-163) or 11.1 mg.mL^-1^ (OXA-405). Enzyme concentration was measured using a NanoDrop spectrophotometer (ThermoFisher) with extinction coefficients calculated using Expasy ProtParam^40^. Final samples were snap-frozen in liquid nitrogen and stored at −80 °C.

### Ligand soaking and crystal structure determination

The structure of unliganded OXA-48 was determined using diffraction data collected from crystals that grew in 0.1 M HEPES pH 7.5, 33% v/v PEG 400, identified from Morpheus MemGold2 sparse matrix screen. OXA-48, OXA-163 and OXA-405 crystals used for structure determination of nacubactam-bound complexes were grown in 0.1 M Tris pH 8.8 – 9.0, 20 – 50% PEG 400 at 10 °C, whilst avibactam-bound and unliganded structures of OXA-163 and OXA-405 were determined from crystals grown in 0.1 M Tris pH 8.5 – 9.0, 28 – 32% PEG 550 at 19 °C. Crystals were soaked with either 5 mM nacubactam or 15 – 100 mM avibactam and either cryoprotected with 20% glycerol before freezing, or frozen directly by immersion in liquid nitrogen. See Table S3 for further details on crystallisation and ligand soaking experiments.

X-ray diffraction data were collected at beamline I03 of Diamond Light Source (Didcot, UK) or PROXIMA 2A beamline of SOLEIL (Paris, France) (see Table S2 for X-ray collection and refinement statistics). Diffraction images were processed using the in-house Xia2 dials and Xia2 3dii pipelines, or merged via AIMLESS (CCP4), ensuring CC_1/2_ greater than 0.3^41,42^. 5% of reflections in each dataset were reserved to calculate *R*_free_ values. Phases for uncomplexed structures were solved by molecular replacement in PhaserMR (Phenix) using an AlphaFold2 prediction as a search model^43,44^. Structures were then refined in phenix.refine using their respective uncomplexed structure (with waters removed) as a starting model for one round of rigid body fitting, followed by manual rebuilding and local refitting in Coot^45,46^. Ligands were fitted into active site *F*_o_-*F*_c_ difference density with ligand geometry restraints generated by Grade2^47^. Figures of structures were generated in open-source PyMOL (Schrödinger)^48^.

### Enzyme kinetics

All enzyme kinetics experiments were performed in 50 mM HEPES pH 7.6, 50 mM NaPO_4_ buffer supplemented with 50 mM NaHCO_3_ and 50 µg.mL^-1^ bovine serum albumin (BSA) in Greiner 96-well half area plates. Nitrocefin hydrolysis was determined with 1 nM enzyme, measuring changing absorbance at 486 nm over 10 mins (Δε 486 = 20,500 M^-1^ cm^-1^) using a Clariostar plate reader (BMG LabTech)^49^. Steady-state parameters (*K*_M_, *k*_cat_) were calculated in Graphpad Prism v10.2 (Table S6). IC_50_ values were determined as described in ^16^, pre-incubating 1 nM enzyme with differing concentrations of inhibitor diluted in kinetics buffer for 10 minutes before adding 75 µM nitrocefin and measuring change in absorbance at 486 nm over 10 mins.

### Minimum inhibitory concentrations (MICs)

A gene encoding OXA-48 was cloned from pSU18 OXA-48^51^ into an empty pUBYT vector^52^. OXA-163 and OXA-405 vectors were then generated from pUBYT OXA-48 using QuikChange Lightning site-directed mutagenesis kit (Agilent) (primers listed in Table S5). *E. coli* TOP10 (lab strain) and ATCC25922 (clinical reference strain) were transformed with their respective plasmids by heat shock. Differing concentrations of β-lactams were combined with 4 µg/mL avibactam (AVI) or nacubactam (NAC), dissolved in DMSO. MICs were determined by broth microdilution method using Mueller-Hinton broth (Sigma), following CLSI guidelines in 96-well microtiter plates (Corning)^53^. The microtiter plate was incubated for 16 – 20 hrs at 37 °C, and absorbance read on a Clariostar plate reader (BMG LabTech) at 600 nm.

### Molecular dynamics simulations

Extended molecular mechanics/molecular dynamics (MM MD) simulations (1.5 µs) were run for the uncomplexed, nacubactam- and avibactam-bound OXA-48, OXA-163 and OXA-405 structures. As a starting point for MM MD simulations a previously determined structure of OXA-48 bound to avibactam (PDB 4S2K^14^) was re-refined in phenix.refine^45^ with a chloride ion added to the dimer interface following inspection of the *F*_o_-*F*_c_ difference map, replacing the water originally modelled. As two dimers reside in the asymmetric unit of 4S2K, chains A and C were removed. Starting structures for simulations of uncomplexed and nacubactam-bound OXA-405 had sections of the β7 - α10 loop that were not modelled in the final crystal structure (residues 237-242) added using Fit Loop (by Rama Search) in Coot^46^. All starting models had crystallographic precipitant molecules removed, excepting the chloride ion found at the dimer interface of all enzyme structures. Amino acid protonation states were altered as predicted by PropKa, based on a system at pH 7.4^29^. Partial charges for avibactam, nacubactam and the carbamylated active site lysine were calculated using the R.E.D webserver^50^. Ligand forcefields were generated using General Amber Force Field (GAFF) parameters and the ff14SB MM forcefield was used for standard protein residues. Hydrogens were added to each complex by tleap (Amber20^51^) and systems solvated within a TIP3P water box^52^ whose edges were at least 10 Å from any protein atoms, with sodium counterions added to balance the overall system charge. All non-water atoms were initially restrained (restraint weight of 100 kcal.mol^-1^.A^-2^) for water minimisation (100 cycles of steepest descent followed by 200 cycles of conjugate gradient), followed by minimisation of all hydrogen atoms, water molecules and chloride ions (1000 cycles of steepest descent, 2000 cycles of conjugate gradient). Systems were then heated to 298 K over 20 ps of simulation using a Langevin thermostat, and system pressure was equilibrated to 1.0 bar over 500 ps with a Berendsen barostat, restraining just the Cα atoms in both steps (restraint weights were 5 kcal.mol^-1^.A^-2^). Production simulations were run over 500 ns, repeating three times for a total sampling time of 1.5 µs for each complex. All simulation analyses were performed using CPPTRAJ^53^, with RMSD values calculated relative to the first deposited simulation structure (at 0.4 ns), excluding all hydrogen atoms and N-terminal residues up to and including Glu24 in both chains of the homodimer.

## Acknowledgements

All simulations were conducted using the facilities of the Advanced Computing Research Centre at the University of Bristol (http://www.bris.ac.uk/acrc/). The authors thank Diamond Light Source (beamline I03, proposals 23269 and 31440) and the SOLEIL synchrotron (beamline PROXIMA 2A), and the respective beamline scientists for their support collecting the X-ray diffraction data presented here. K.E.G., J.S. and C.J.S. acknowledge funding from the U.K. Biotechnology and Biological Sciences Research Council (BBSRC, BB/W001187/1). M.B. was supported by the BBSRC-funded South West Biosciences Doctoral Training Partnership (BB/T008741/10). This work was supported in part by grant MR/W006308/1 for the GW4 BIOMED2 DTP, awarded to the Universities of Bath, Bristol, Cardiff and Exeter from the Medical Research Council (MRC)/UKRI. C.L.T., J.S. and A.J.M. thank the Medical Research Council for support through the grant MR/T016035/1. This work is part of a project that has received funding from the European Research Council under the European Horizon 2020 research and innovation program (PREDACTED Advanced Grant Agreement no. 101021207) to A.J.M. and J.S.

## Abbreviations

BLI: β-lactamase inhibitor
DBO: diazabicyclooctane
MM MD: molecular mechanics molecular dynamics
OXA: oxacillinase
RMSD: root mean-squared deviation
RMSF: root mean-squared fluctuation
SBL: serine β-lactamase

## Supporting Information

Sequences of oligonucleotide primers; conditions, data processing and validation statistics for crystallographic experiments; steady-state kinetic parameters for hydrolysis of reporter substrate (nitrocefin); supplementary images of crystal structures; analyses of molecular simulations (DOC)

## Author Contributions

J.F.H., K.E.G., C.J.S. and J.S. conceived the experiments. J.F.H. and K.E.G. performed laboratory experiments and crystallographic data collection/processing, with supervision from C.L.T., P.H, J.M.S and Y.T. J.F.H. undertook molecular simulations with training and supervision from M.B. J.F.H. and K.E.G. drafted the manuscript, with revisions by J.S. and input from and final approval by all authors.

## Data availability statement

Structure factors and coordinates for all crystal structures presented here have been deposited with the Protein Data Bank (PDB) with the following accession numbers: uncomplexed OXA-48 (9H11), OXA-48 in complex with nacubactam (9H12), uncomplexed OXA-163 (9H13), OXA-163 in complex with avibactam (9H14), OXA- 163 in complex with nacubactam (9H15), uncomplexed OXA-405 (9H16), OXA-405 in complex with avibactam (9H17), OXA-405 in complex with nacubactam (9H18). All raw MD simulations data (including equilibrium and non-equilibrium simulations) will be made freely available at the University of Bristol Research Data Repository (https://data.bris.ac.uk/). Analysis scripts will be made available upon request.

