## Supplementary material for "Electrostatic interactions define nacubactam potency against OXA-48-like β-lactamases": OXA48_nacubactam_paper_SI_BioRxiv.pdf

Supporting Information

**Supporting Information**

Contents

Table S1: Minimum inhibitory concentrations (MICs) of β-lactams combined with DBO inhibitors against E. coli

Table S2: X-ray data collection and structure refinement statistics

Table S3: Crystallisation and ligand soaking experiment summary

Table S4: DBO complex ligand and KCX73 fitting statistics

Table S5: Primer sequences used to create OXA-163 and OXA-405 expression and MIC constructs

Table S6: Steady state kinetic parameters for nitrocefin hydrolysis by OXA-48, OXA-163 and OXA-405

Figure S1: Active arrangement site of uncomplexed OXA-48

Figure S2: *In crystallo* conformation of Arg214 in uncomplexed and DBO-bound OXA-48

Figure S3: Nacubactam ionisable group predicted pKa values

Figure S4: RMSD plots of MM MD simulation trajectories

Figure S5: Distance analysis of Arg214 relative to avibactam and nacubactam across MD simulation trajectories

Figure S6: OXA-48 deacylating water channel rotamer analysis over MD simulations

Figure S7: DBO-derived carbamoyl-enzyme unbiased Fo-Fc omit maps

Figure S8: Overlay of OXA-48, OXA-163 and OXA-405 apoenzyme crystal structures

Figure S9: Tyr211:β7 – α10 loop interactions in the OXA-405:nacubactam complex

| | β-Lactam | Ertapenem | | | Meropenem | | | Ceftazidime | | | - | |
|---|---|---|---|---|---|---|---|---|---|---|---|---|
| | **Inhibitor** | - | +AVI | +NAC | - | +AVI | +NAC | - | +AVI | +NAC | +AVI | +NAC |
| **Strain / Plasmid** | *E. coli* TOP10 pUBYT OXA-48 | 64 | 1 | <0.25 | 8 - 16 | 1 - 2 | 8 | 4 | 4 | <0.125 | >32 | >64 |
| | *E. coli* TOP10 pUBYT OXA-163 | 8 | 2 | <0.016 | 1 | 0.5 | 0.5 - 1 | >128 | 8 - 16 | 0.5 | >32 | >64 |
| | *E. coli* TOP10 pUBYT OXA-405 | 8 | 1 | <0.016 | 1 -2 | 0.5 | 1 | >128 | 8 | 0.5 | >32 | >64 |
| | *E. coli* TOP10 pUBYT | 0.25 | 0.25 | <0.016 | 0.25 | 0.25 | <0.016 | 2 - 4 | 2 - 4 | 0.25 | >32 | >64 |
| | *E. coli* ATCC25922 | 0.125 - 0.25 | 0.125 | <0.016 | 0.125 - 0.25 | 0.125 | <0.016 | 4 | 2 | 0.125 | >32 | >64 |

**Table S1: Minimum inhibitory concentrations (MICs) of β-lactams combined with DBO inhibitors against *E. coli*.**

| | OXA-48 uncomplexed[a] | OXA48: nacubactam[a] | OXA-163 uncomplexed[b] | OXA163: avibactam[a] | OXA-163: nacubactam[a] | OXA-405 uncomplexed[a] | OXA-405: avibactam[a] | OXA-405: nacubactam[a] |
|---|---|---|---|---|---|---|---|---|
| **PDB ID** | 9H11 | 9H12 | 9H13 | 9H14 | 9H15 | 9H16 | 9H17 | 9H18 |
| **Data collection** | | | | | | | | |
| Space group | $P6_522$ | $P6_522$ | $P6_522$ | $P6_522$ | $P2_12_12$ | $P6_522$ | $P6_522$ | $P6_522$ |
| Molecules/ASU | 2 | 2 | 2 | 2 | 2 | 2 | 2 | 2 |
| *Cell dimensions* | | | | | | | | |
| a, b, c (Å) | 123.51, 123.51, 161.56 | 121.76, 121.76, 159.84 | 122.55, 122.55, 160.43 | 122.60, 122.60, 160.53 | 70.62, 79.04, 126.25 | 122.08, 122.08, 159.12 | 122.30, 122.30, 160.18 | 123.69, 123.69, 159.35 |
| α, β, γ (°) | 90, 90, 120 | 90, 90, 120 | 90, 90, 120 | 90, 90, 90 | 90, 90, 90 | 90, 90, 120 | 90, 90, 120 | 90, 90, 120 |
| Wavelength (Å) | 0.8622 | 0.9763 | 0.9000 | 0.9500 | 0.9763 | 0.9795 | 0.9537 | 0.9763 |
| Resolution (Å) | 61.76 - 1.38 (1.40 - 1.38) | 60.88 - 1.51 (1.54 - 1.51) | 48.69 - 1.44 (1.46 - 1.44) | 106.17 - 1.56 (1.59 - 1.56) | 52.66 - 2.18 (2.22 - 2.18) | 63.57 - 1.56 (1.59 - 1.56) | 57.13 - 1.46 (1.49 - 1.46) | 57.66 - 1.33 (1.35 - 1.33) |
| $R_{pim}$ ($I$) | 0.055 (1.020) | 0.047 (0.956) | 0.030 (1.665) | 0.031 (0.899) | 0.141 (1.171) | 0.053 (1.629) | 0.030 (0.818) | 0.027 (1.319) |
| $CC_{1/2}$ | 0.999 (0.309) | 0.999 (0.327) | 0.999 (0.348) | 0.999 (0.313) | 0.987 (0.327) | 0.997 (0.348) | 0.999 (0.286) | 0.999 (0.297) |
| $I/\sigma$ ($I$) | 8.9 (0.8) | 11.3 (0.8) | 13.5 (0.4) | 12.8 (0.9) | 5.1 (0.6) | 8.0 (0.5) | 12.0 (0.5) | 8.9 (0.1) |
| Completeness (%) | 100 (100) | 100 (100) | 99.9 (99.7) | 100 (100) | 98.7 (98.1) | 100 (100) | 100 (100) | 100 (98.1) |
| Redundancy | 40.2 (41.1) | 39.5 (33.5) | 40.1 (38.9) | 38.7 (39.1) | 14.1 (14.6) | 13.4 (13.3) | 41.8 (42.2) | 39.9 (40.5) |
| **Refinement** | | | | | | | | |
| Resolution (Å) | 61.76 - 1.38 (1.43 - 1.38) | 60.88 - 1.51 (1.56 - 1.51) | 48.69 - 1.44 (1.49 - 1.44) | 64.03 - 1.56 (1.62 - 1.56) | 52.66 - 2.18 (2.26 - 2.18) | 61.04 - 1.56 (1.62 - 1.56) | 57.13 - 1.46 (1.51 - 1.46) | 57.66 - 1.33 (1.38 - 1.33) |
| No. reflections | 148,214 | 109,282 | 127,598 | 101,080 | 37,053 | 99,172 | 121,931 | 138,681 |
| $R_{work}$ / $R_{free}$ | 0.170 / 0.187 | 0.173 / 0.192 | 0.168 / 0.186 | 0.164 / 0.185 | 0.196 / 0.243 | 0.175 / 0.193 | 0.175 / 0.197 | 0.182 / 0.199 |
| *No. atoms* | | | | | | | | |
| Protein | 4104 | 4116 | 4104 | 4052 | 3947 | 4025 | 4003 | 4031 |
| Solvent | 688 | 506 | 568 | 436 | 245 | 575 | 458 | 595 |
| Ligand | - | 63 | - | 34 | 42 | - | 34 | 42 |
| *B-factors* (Å$^2$) | | | | | | | | |
| Protein | 17.7 | 20.4 | 25.3 | 29.2 | 40.3 | 27.7 | 25.6 | 28.0 |
| Solvent | 30.3 | 34.6 | 38.4 | 39.2 | 44.2 | 39.7 | 37.0 | 35.4 |
| Ligand | - | 20.8 | - | 38.7 | 44.4 | - | 30.6 | 34.01 |
| *RMS deviations* | | | | | | | | |
| Bond angles (°) | 0.78 | 0.84 | 0.79 | 0.82 | 0.84 | 0.84 | 0.82 | 0.81 |
| Bond lengths (Å) | 0.0054 | 0.0064 | 0.0057 | 0.0064 | 0.0073 | 0.0064 | 0.0058 | 0.0062 |
| *Ramachandran (%)* | | | | | | | | |
| Outliers | 0 | 0 | 0 | 0 | 0 | 0 | 0.21 | 0 |
| Favoured | 97.92 | 98.56 | 97.03 | 97.49 | 96.84 | 97.17 | 97.68 | 96.82 |

**Table S2: X-ray data collection and structure refinement statistics.** [a]*Experiments performed at Diamond Light Source (I03 beamline, Didcot, UK) or* [b]*SOLEIL (PROXIMA 2A beamline, Paris, France). Outer shell statistics are shown in brackets.*

| Protein | Ligand | Ligand soak time | Ligand concentration | Cryoprotection | Crystallisation condition | Temp (°C) | Seed |
|---|---|---|---|---|---|---|---|
| OXA-48 | Apoenzyme | - | - | - | 0.1 M HEPES pH 7.5, 33% PEG 400 | 10 | - |
| | Nacubactam | 30 mins | 5 mM | 20% Glycerol | 0.1 M Tris pH 8.8, 50% PEG 400 | 10 | - |
| OXA-163 | Apoenzyme | - | - | - | 0.1 M Tris pH 9.0, 32% PEG 550 | 19 | OXA-163 crystals |
| | Nacubactam | 4 hour | 5 mM | 20% Glycerol | 0.1 M Tris pH 8.8, 40% PEG 400 | 10 | OXA-163 crystals |
| | Avibactam | 1 hour | 15 mM | | 0.1 M Tris pH 9.0, 32% PEG 550 | 19 | OXA-163 crystals |
| OXA-405 | Apoenzyme | - | - | - | 0.1 M Tris pH 8.5, 28% PEG 550 | 19 | OXA-405 crystals |
| | Nacubactam | 2 hour | 5 mM | - | 0.1 M Tris pH 9.0, 20% PEG 400 | 10 | - |
| | Avibactam | 1 hour | 100 mM | | 0.1 M Tris pH 8.5, 28% PEG 550 | 19 | OXA-405 crystals |

**Table S3: Crystallisation and ligand soaking experiment summary.** *Seed was generated using Seed Bead kit and Crystal Crusher (Hampton Research).*

**A.**

| **Structure** | **KCX73 occupancy** | **KCX73 RSCC** |
|---|---|---|
| OXA-48:nacubactam | - | - |
| OXA-163:avibactam | 0.28 / - | 0.97 / - |
| OXA-163:nacubactam | - | - |
| OXA-405:avibactam | 0.41 / 0.31 | 0.97 / 0.96 |
| OXA-405:nacubactam | - | - |

**B.**

| **Structure** | **Ligand occupancy** | **Ligand RSCC** |
|---|---|---|
| OXA-48:nacubactam | 0.62, 0.38 / 1 | 0.97, 0.97 / 0.96 |
| OXA-163:avibactam | 1 / 1 | 0.91 / 0.87 |
| OXA-163:nacubactam | 1 / 1 | 0.94 / 0.95 |
| OXA-405:avibactam | 1 / 1 | 0.90 / 0.93 |
| OXA-405:nacubactam | 1 / 1 | 0.92 / 0.94 |

**Table S4: DBO complex ligand and KCX73 fitting statistics.** *Statistics calculated by PDB validation server.*

| Reaction | Forward primer | Reverse primer |
|---|---|---|
| pOPIN-F OXA-163 cloning | 5’-AAGTTCTGTTTCAGGGCCCGAAAGAATGGCAGGAAAACAAGAGCTG-3’ | 5’-ATGGTCTAGAAAGCTTTACGGGATGATTTTCTCCTGTTTGAG-3’ |
| pOPIN-F OXA-405 cloning | 5’-AAGTTCTGTTTCAGGGCCCGAAAGAATGGCAGGAAAACAAATCCTG-3’ | 5’-ATGGTCTAGAAAGCTTTACGGGATGATCTTTTCCTGCTTCAAC-3’ |
| pUBYT OXA-48 to OXA-163 214-217 deletion | 5’-AAAACTGGATACTCGACTAAGATTGGCTGGTGGGTC-3’ | 5’-GACCCACCAGCCAATCTTAGTCGAGTATCCAGTTTT-3’ |
| pUBYT OXA-48 to OXA-163 S212D substitution | 5’-CGGGCTAAAACTGGATACGCTACTAAGATTGGCTGGTGG-3’ | 5’-CCACCAGCCAATCTTAGTAGCGTATCCAGTTTTAGCCCG-3’ |
| pUBYT OXA-405 214-217 deletion | 5’-GCTAAAACTGGATACTCGCCTAAGATTGGCTGGTGG-3’ | 5’-CCACCAGCCAATCTTAGGCGAGTATCCAGTTTTAGC-3’ |

**Table S5: Primer sequences used to create OXA-163 and OXA-405 expression and MIC constructs.**

| Enzyme | $K_M$ (µM) | $k_{cat}$ ($s^{-1}$) | $k_{cat}/K_M$ ($s^{-1}$ $\mu M^{-1}$) |
|---|---|---|---|
| **OXA-48** | 162.5 | 581.1 | 3.6 |
| **OXA-163** | 14.5 | 57.6 | 4.0 |
| **OXA-405** | 21.6 | 99.7 | 4.6 |

**Table S6: Steady state kinetic parameters for nitrocefin hydrolysis by OXA-48, OXA-163 and OXA-405.**

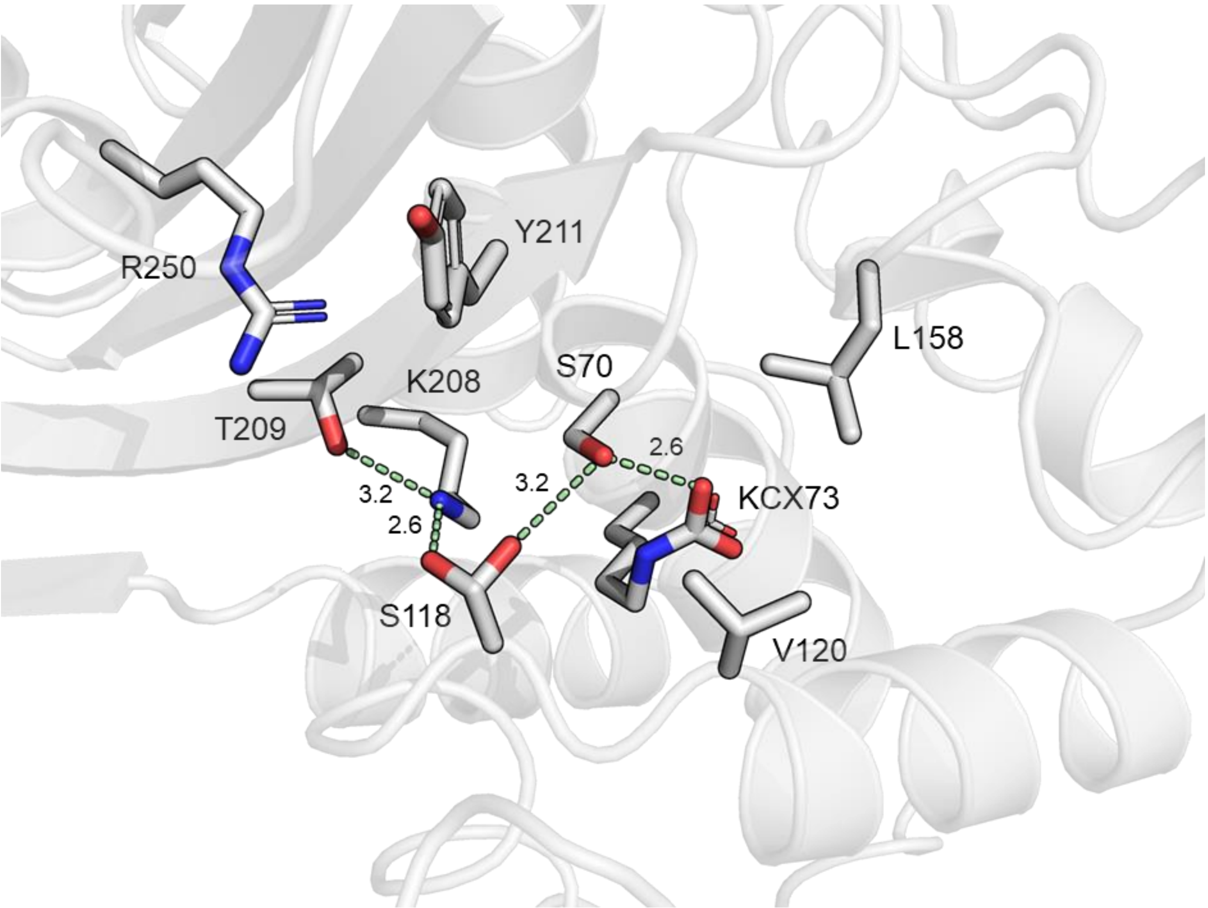


**Figure S1: Active arrangement site of uncomplexed OXA-48.** *Side-chains of key chain A active site residues are shown as grey sticks, with hydrogen bonds between them labelled as pale green dashes, with distances labelled in*

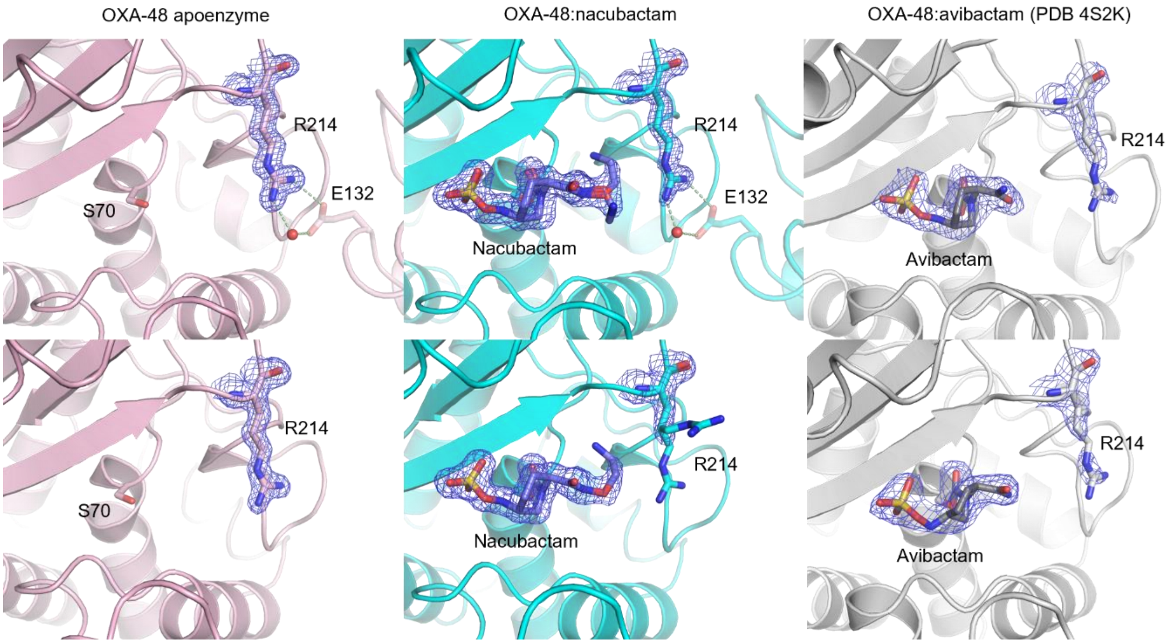


**Figure S2: *In crystallo* conformation of Arg214 in uncomplexed and DBO-bound OXA-48.** *Arg214 and DBO final* 2$F_o$-$F_c$ *electron density, contoured to 1σ. Interacting crystal symmetry subunits not in the asymmetric unit are shown as transparent sticks/cartoon and Arg214-mediated crystal packing interactions highlighted by dashed lines, with bridging waters shown as red spheres.*

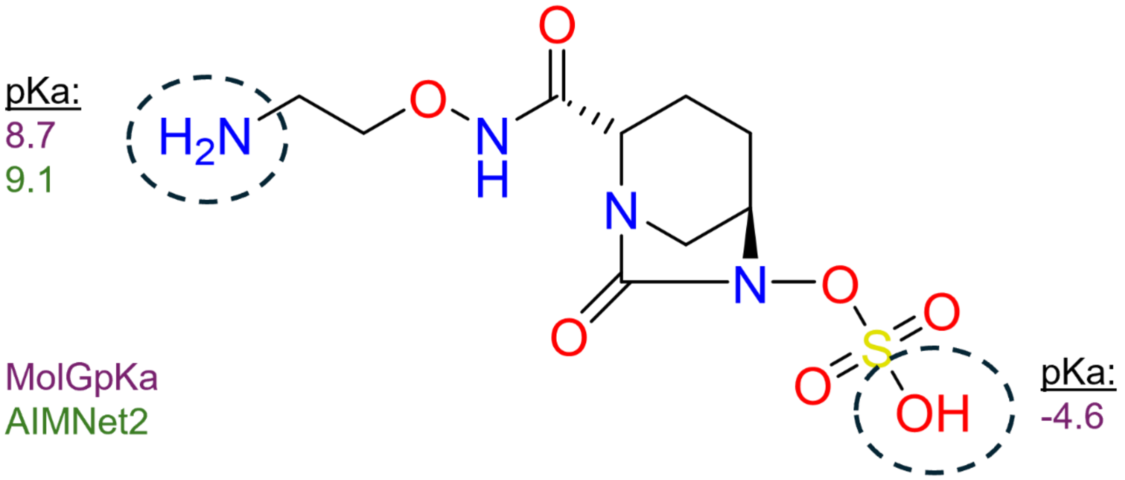


**Figure S3: Nacubactam ionisable group predicted pKa values.** *MolGpKa*[1] *and AIMNet2*[2] *webserver pKa predictions are shown adjacent to ionisable groups of nacubactam, highlighted by a dashed circle.*

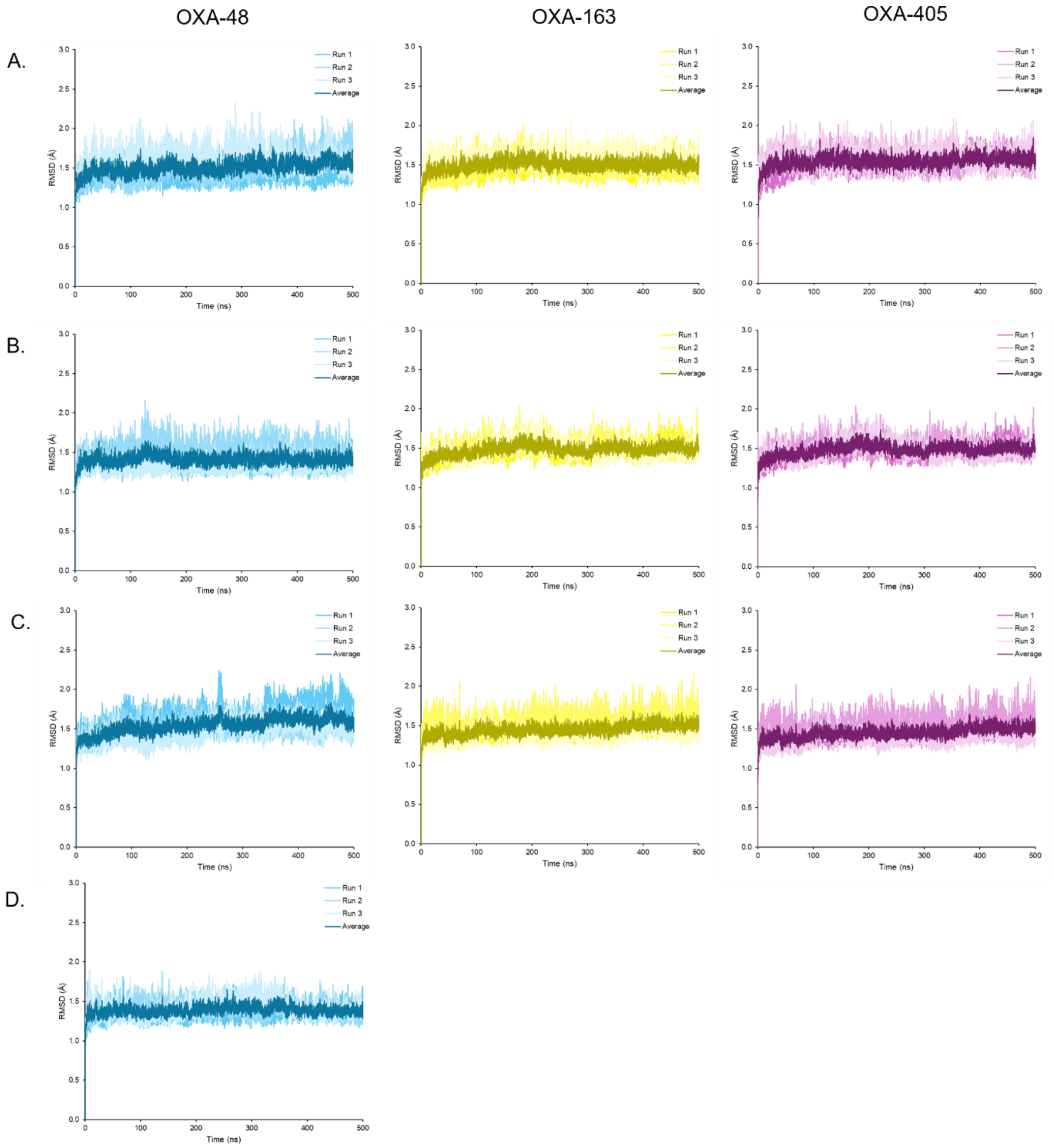


**Figure S4: RMSD plots of MM MD simulation trajectories.** *Simulations of (A.) uncomplexed, (B.) avibactam-bound, (C.) nacubactam-bound (C2 tail N protonated) and (D.) nacubactam-bound (C2 tail N deprotonated) OXA-48 (blue), OXA-163 (yellow), OXA-405 (pink) complexes. Each 500 ns simulation run is shown separately, with the average RMSD shown in the darkest shade of their respective colours.*

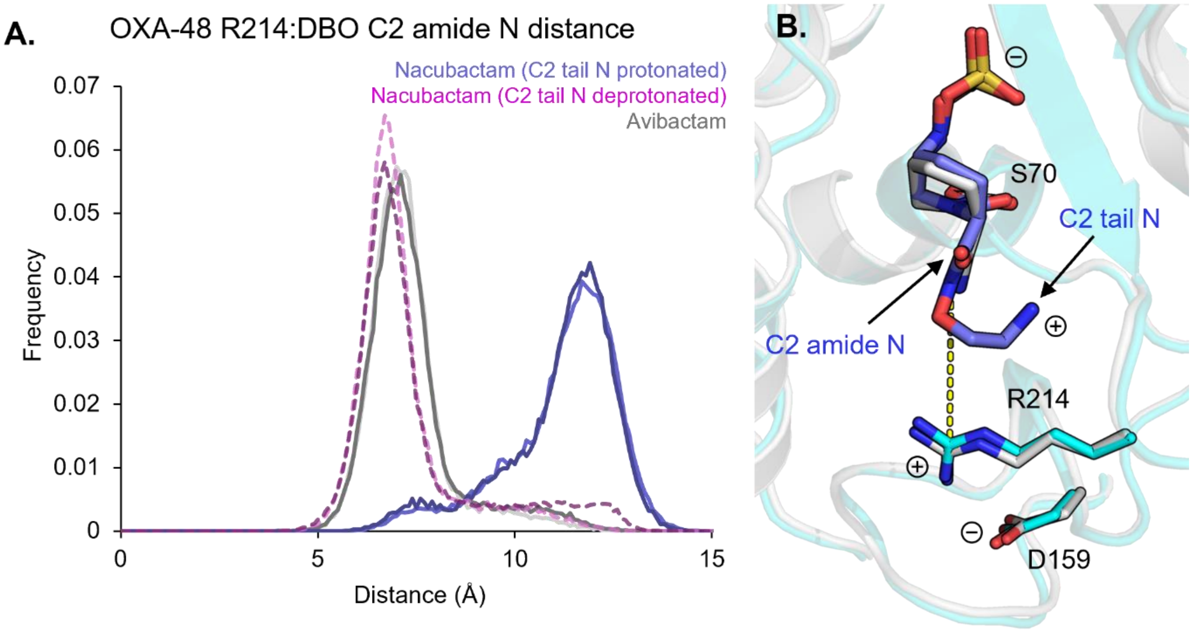


**Figure S5: Distance analysis of Arg214 relative to avibactam and nacubactam across MD simulation trajectories.** *(A) Frequency histogram of Arg214-C2 amide nitrogen distance over the OXA-48:DBO MD simulations. The darker shades of each colour correspond to the chain B active site complexes with OXA-48 and the lighter for chain A. (B) OXA-48 active site with avibactam and nacubactam overlayed, the distance between Arg214 and the DBO C2 amide nitrogen measured in panel A is labelled as a yellow dashed line.*

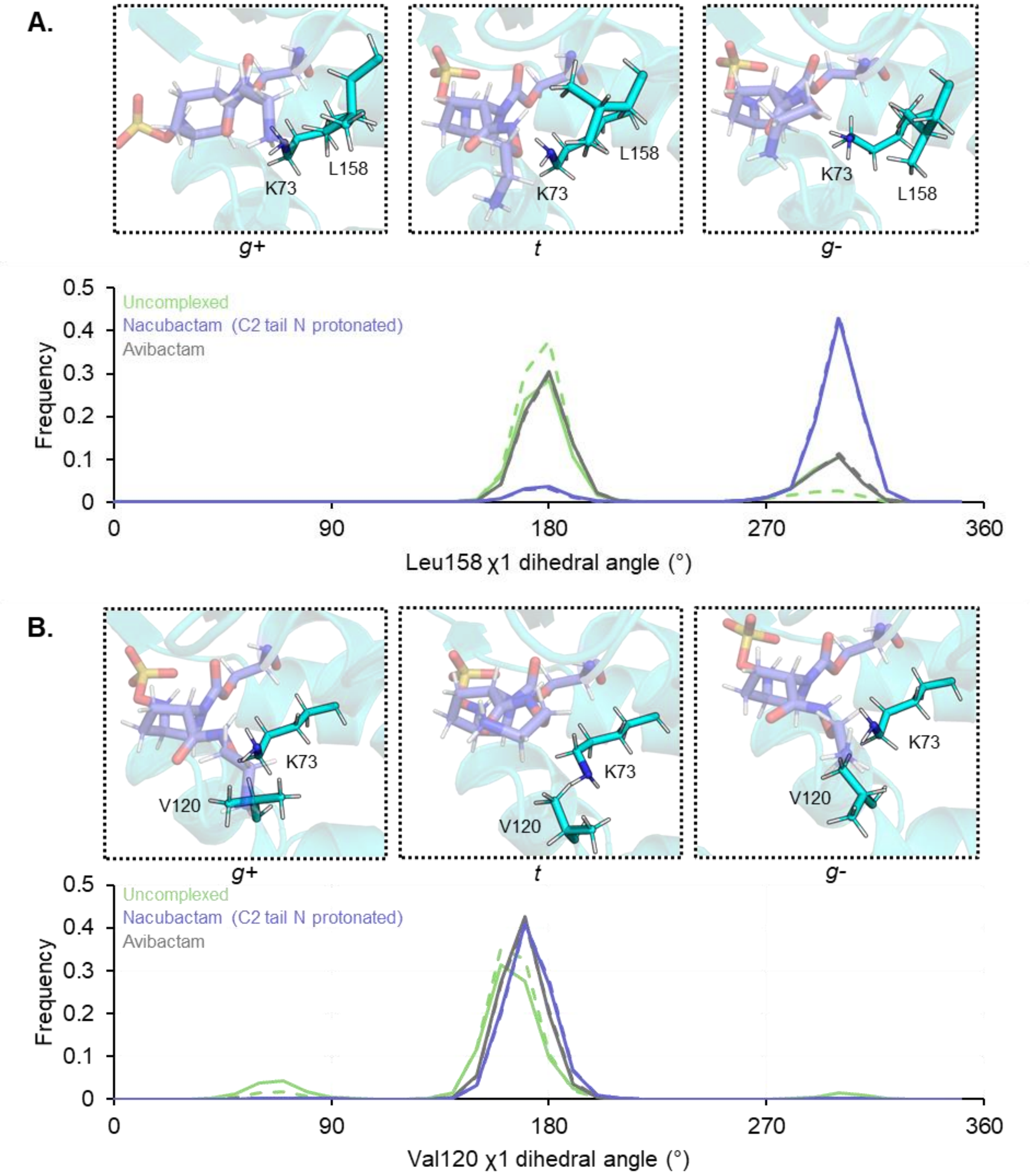


**Figure S6: OXA-48 deacylating water channel rotamer analysis over MD simulations.** *(A) Leu158 $\chi_1$ dihedral angle (N-Cα-Cβ-Cγ) and (B) Val120 $\chi_1$ dihedral angle (N-Cα-Cβ-Cγ1) frequency histograms for simulation trajectories of nacubactam-bound (blue), avibactam-bound (grey) and uncomplexed (green) OXA-48.*

*Representative simulation snapshots are shown above in the* g+ *(~60°),* t *(~180°) and* g- *(~300°)* $\chi_1$ *rotamers of Leu158 and Val120. Lys73 and nacubactam carbamoyl-enzyme complex (blue, transparent) are also shown as sticks.*

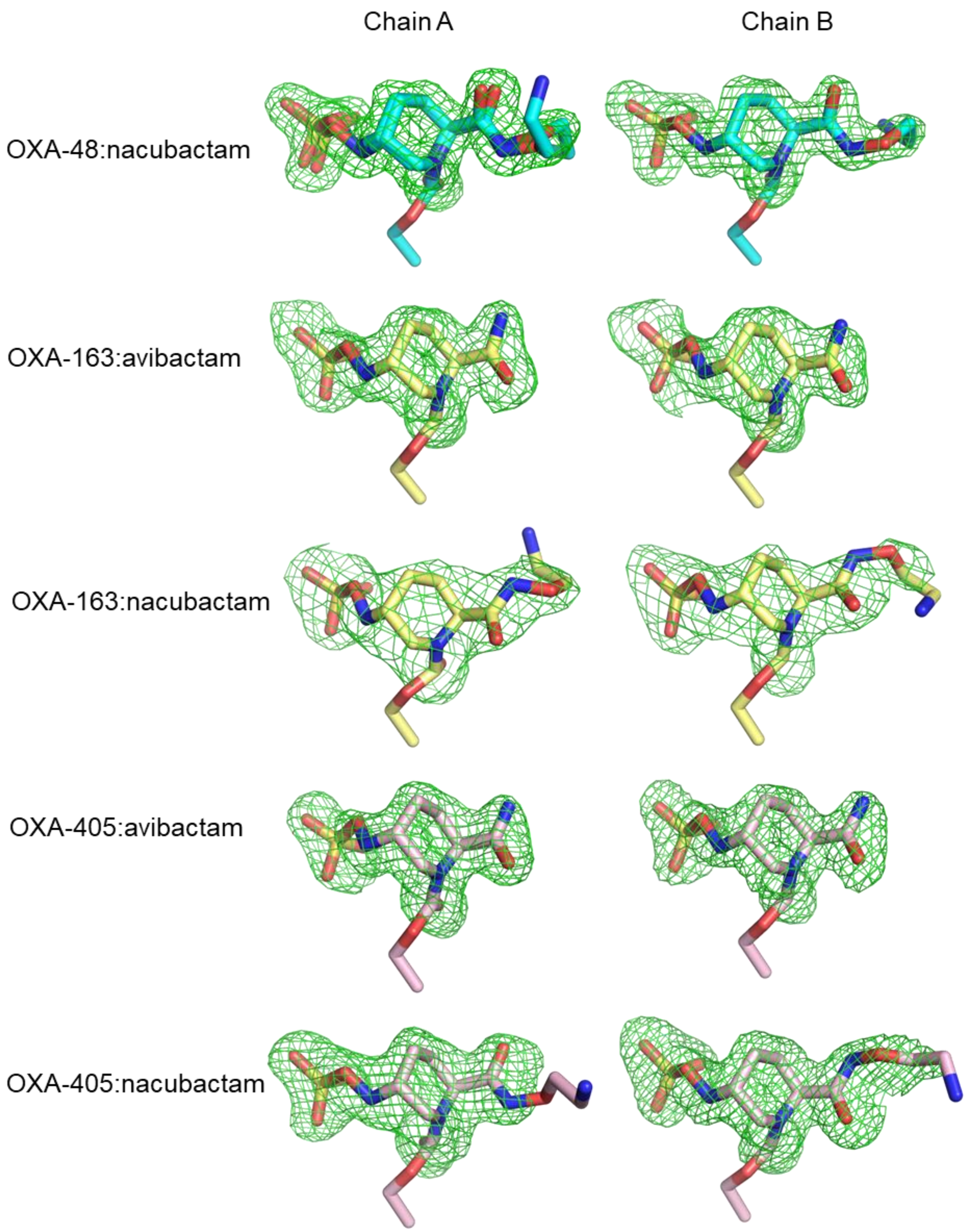


**Figure S7: DBO-derived carbamoyl-enzyme unbiased Fo-Fc omit maps.** *Maps contoured to 3σ (green mesh). DBO-Ser70 active site carbamoyl-enzyme complexes are shown as sticks*

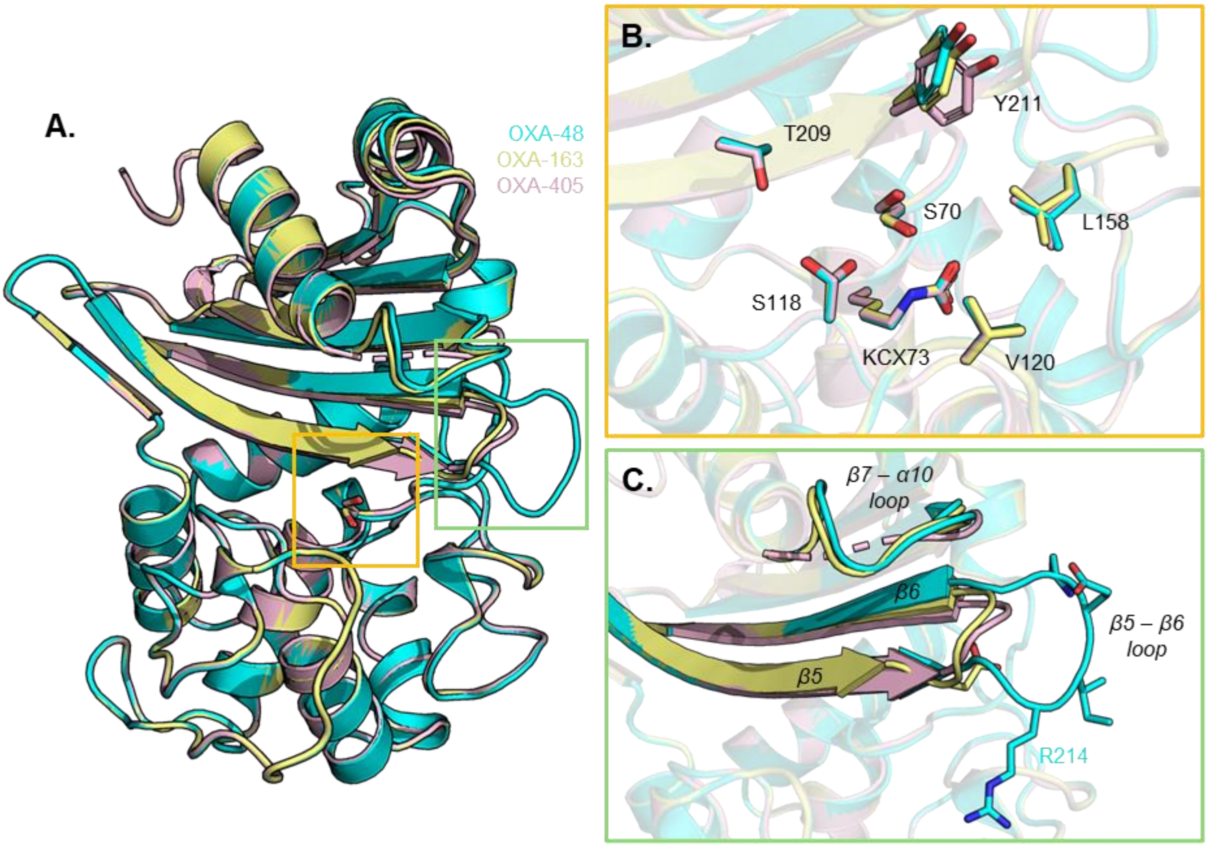


**Figure S8: Overlay of OXA-48, OXA-163 and OXA-405 apoenzyme crystal structures.** *(A) Chain B overall topology, with zoom ins of the (A) active site and (B) β5 – β6 loops. RMSD values of the overlay are 0.117 Å and 0.141 Å for OXA-163 and OXA-405 apoenzymes respectively, relative to OXA-48. Key active site residues and substituted/deleted residues across OXA-48 variants are shown as sticks. Unmodelled residues are represented by a dashed line.*

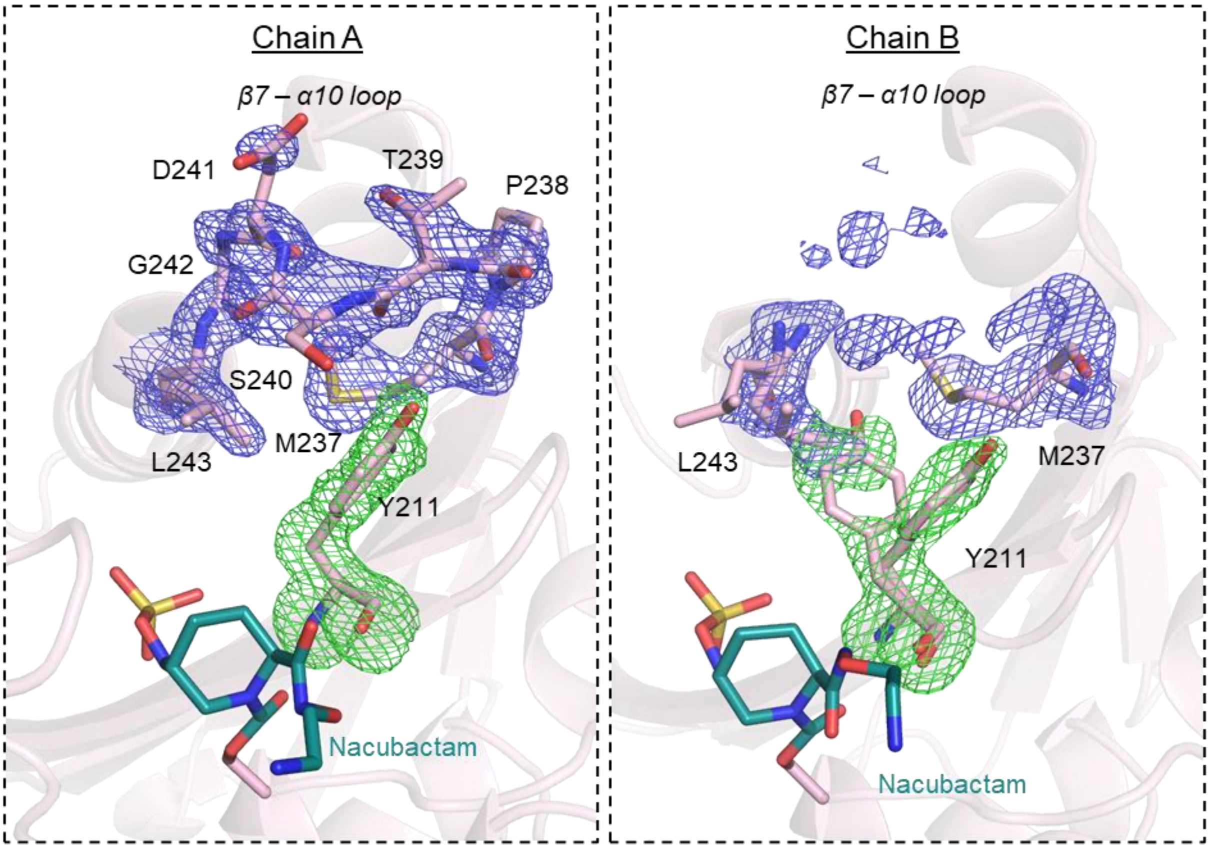


**Figure S9: Tyr211:β7 – α10 loop interactions in the OXA-405:nacubactam complex.** *Tyr211 unbiased* $F_o$-$F_c$ *omit map is shown as a green mesh, contoured to 3σ. The β7 – α10 (residues 237-243) final* $2F_o$-$F_c$ *electron density is represented as a blue mesh, contoured to 1σ. The chain B β7 – α10 map* $2F_o$-$F_c$ *was generated using the chain A loop superimposed onto the chain B of OXA-405, and contouring the electron density around the overlaid loop.*
